# Analysis of single-cell CRISPR perturbations indicates that enhancers act multiplicatively and provides limited evidence for epistatic-like interactions

**DOI:** 10.1101/2023.04.26.538501

**Authors:** Jessica Zhou, Karthik Guruvayurappan, Shushan Toneyan, Hsiuyi V. Chen, Aaron R. Chen, Peter Koo, Graham McVicker

## Abstract

A single gene may have multiple enhancers, but how they work in concert to regulate transcription is poorly understood. To analyze enhancer interactions throughout the genome, we developed a generalized linear modeling framework, GLiMMIRS, for interrogating enhancer effects from single-cell CRISPR experiments. We applied GLiMMIRS to a published dataset and tested for interactions between 46,166 enhancer pairs and corresponding genes, including 264 ’high-confidence’ enhancer pairs. We found that enhancer effects combine multiplicatively but with limited evidence for further interactions. Only 31 enhancer pairs exhibited significant interactions (FDR < 0.1), of which none came from the high confidence subset and 20 were driven by outlier expression values. Additional analyses of a second CRISPR dataset and in silico enhancer perturbations with Enformer both support a multiplicative model of enhancer effects without interactions. Altogether, our results indicate that enhancer interactions are uncommon or have small effects that are difficult to detect.

**Highlights:**

- Analysis of a large single-cell CRISPRi screen finds limited evidence for synergistic or redundant interactions between enhancers
- The collective action of multiple enhancers on a common target gene follows a multiplicative model of activity
- A new statistical framework for simulating and modeling data from single-cell CRISPRi screens

## Introduction

Enhancers are distal cis-regulatory elements that direct transcription and shape cellular identity, growth, and biological function. Most genes are regulated by multiple enhancers^1,2^, yet we lack a detailed understanding of how they act together to influence gene expression. When multiple enhancers for a gene are active in the same cell type, it is often assumed that they act additively—that is, their combined effect is equal to the sum of their individual effects^3^. However, enhancers may also act non-additively, and interactions between regulatory elements may modulate their effects on gene expression^3–10^.

To date, most studies of regulatory elements have examined their effects independently, and studies of regulatory element interactions have focused on a small number of loci and reached contradictory conclusions^4–8^. For example, a study of *ɑ-globin* regulation in mice found that this gene’s expression is best explained by simple additivity between constituent elements of its super enhancer^7^. In addition, a study that systematically deleted three constituent enhancers of a super enhancer for *Wap* in mice found differences in the magnitudes of effect that each enhancer had on the target gene and no evidence of synergy between the enhancers, with all three enhancers necessary to induce full induction of the gene during pregnancy^8^. Reexamination of both of these super enhancer datasets under hypothetical additive, multiplicative, and logistic activity models found that the effects of the constituent enhancers on the target genes were best described by a logistic model, where enhancers work together multiplicatively until a saturation expression level is reached, but no significant evidence for interactions between enhancers^5^. Contrary to these findings, a recent study of the *MYC* locus described both synergistic and additive enhancer-enhancer interactions, where enhancers separated from one another by larger genomic distances are more likely to have synergistic interactions and enhancers located closer to one another are more likely to have additive interactions^9^. Altogether, these studies have been limited to the examination of a small number of genes and enhancers and their results are difficult to interpret due to their conflicting findings and the lack of explicit definitions and consistent terminology for different models of enhancer activity.

CRISPR-induced perturbations of enhancer sequences have enabled high-throughput quantification of the effects of enhancers on gene expression^11,12^. These experiments have revealed that an activity-by-contact (ABC) model that combines enhancer activity with promoter contact frequency can predict the effect of enhancers on gene expression^11^. However, the ABC model scores each enhancer individually and was not used to predict how joint perturbations to multiple enhancers affect gene expression^11^.

CRISPR perturbations can be combined with single-cell RNA sequencing^10,13–19^ to induce multiple genomic perturbations and measure their effects on gene expression in individual cells. These experiments can be used to identify interactions, or epistatic-like effects, between targeted sequences. Here, we harness this feature of datasets generated by these experiments to measure the combined effects of multiple regulatory elements on gene expression. We present GLiMMIRS (Generalized Linear Models for Measuring Interactions between Regulatory Sequences), a statistical analysis framework that can be applied to single cell CRISPR perturbation experiments to quantify the effects of multiple regulatory elements on gene expression and identify interactions between them. GLiMMIRS has both data simulation and modeling components and can account for variation in gRNA efficiency. We applied GLiMMIRS to a multiplexed, single-cell CRISPR interference (CRISPRi) experiment that targeted putative enhancers in K562 cells^13^ and conducted a power analysis, which found that this dataset provides sufficient power to detect strong interactions between enhancers, and moderate power to detect weak interactions. We also analyzed a second CRISPRi dataset and performed in silico perturbations to enhancer pairs with Enformer, a deep neural network that predicts gene expression from genomic sequences. All three analyses support a model in which enhancers act multiplicatively to affect the expression of their target genes, but provided limited evidence for the presence of additional interactions between them.

## Results

### Detecting Enhancer Effects with GLiMMIRS-base

To analyze the effect of multiple enhancers on gene expression, we leveraged data from a multiplexed, single-cell CRISPRi screen performed in K562 cells^13^. In this screen, which we refer to as Gasperini et al., gRNAs were designed to target putative enhancers and enhancer-gene pairs were identified by associating perturbed enhancers with differences in the expression of nearby genes. Due to the high multiplicity of infection (MOI) used in this experiment, many gRNAs targeting different enhancers are present within each cell. We realized that this feature of the dataset could be leveraged to quantify how pairs of enhancers regulate the expression of common target genes and to detect potential interaction effects between them.

To estimate the effects of regulatory elements on target genes using data from this single-cell CRISPRi screen, we developed GLiMMIRS, a dual modeling and simulation framework. GLiMMIRS consists of three components: GLiMMIRS-base, a baseline model for estimating the regulatory effects of a single enhancer on a target gene (Fig. 1A); GLiMMIRS-int, an interaction model for estimating the combined regulatory effects of two enhancers on a target gene (Fig. 1B); and GLiMMIRS-sim, a simulation framework for single-cell CRISPRi screens (Fig. 1C).

**Figure 1.**
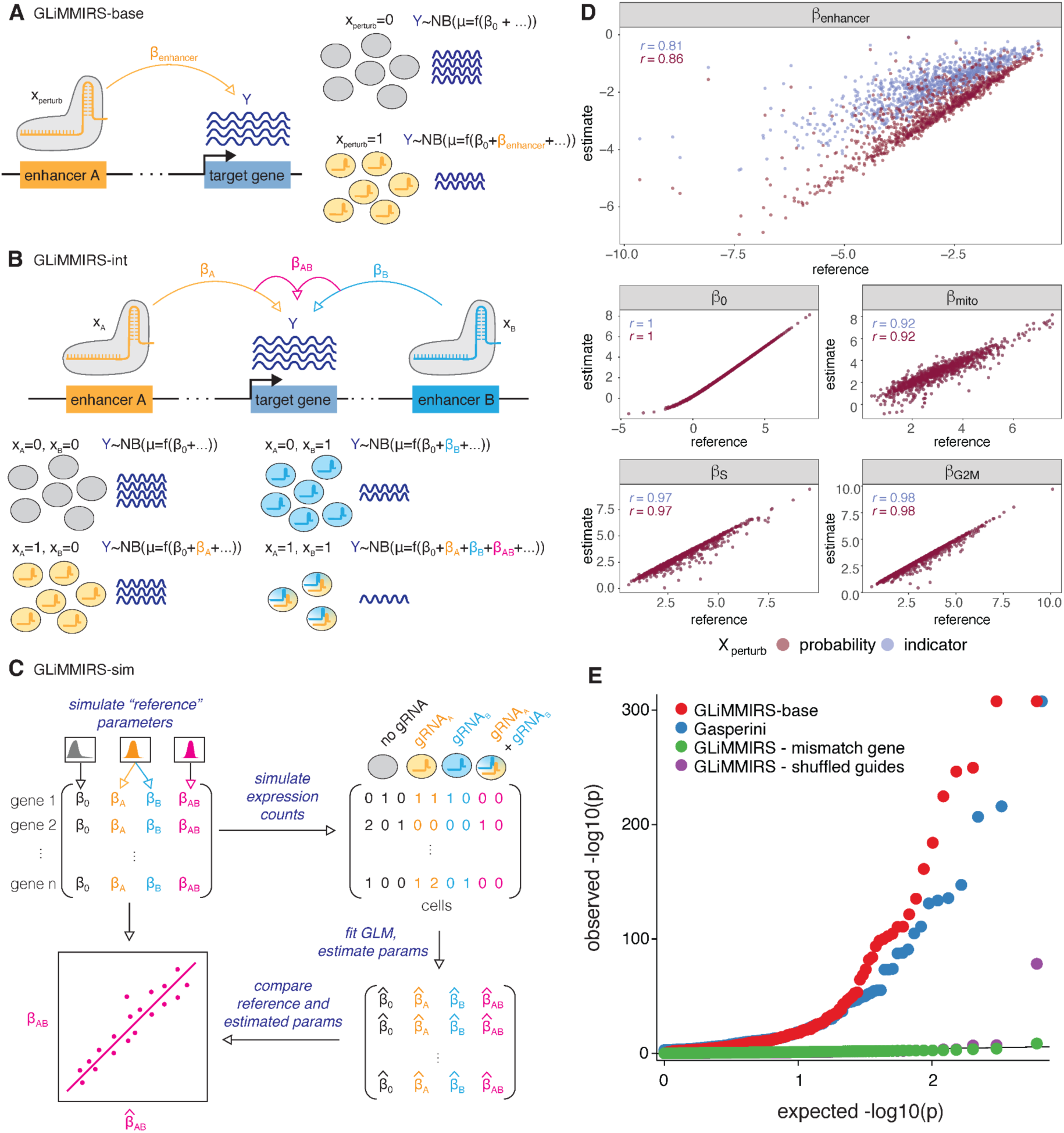
Detecting enhancer effects with GLiMMIRS. **A)** Schematic of GLiMMIRS-base, a GLM to infer the effect of individual enhancers on the expression of target genes. **B)** Schematic of GLiMMIRS-int, a GLM to infer joint perturbation effects of two enhancers on target genes and for detecting interaction effects between the enhancers. **C)** Schematic of GLiMMIRS-sim, which simulates data from single-cell CRISPR perturbation experiments with RNA-seq readout. It can be used to generate reference values for evaluating the performance of GLiMMIRS-base and GLiMMIRS-int. **D)** Scatter plots comparing reference coefficient values generated by GLiMMIRS-sim to coefficient estimates from applying GLiMMIRS-base to the simulated data (y-axis) using two different values of *X_perturb_*: (1) a perturbation probability (magenta), calculated as a function of gRNA efficiencies; and (2) an indicator variable based on the presence/absence of a targeting gRNA for the putative enhancer of interest (lavender). Pearson correlations (r) are denoted on plots (see also Table S2). **E)** Quantile-quantile plot of observed versus expected -log_10_p values. The baseline values (red) are the results from GLiMMIRS-base. The Gasperini values (blue) are previously published results. Mismatch gene (green) is a negative control in which GLiMMIRS-base was applied to randomly selected genes, rather than genes near to the enhancer. Shuffled guides (purple) is a negative control in which the vector containing guide perturbation probabilities for each cell was shuffled.

We first developed GLiMMIRS-base, which is a negative binomial generalized linear model (GLM) that can be fit to single-cell RNA-seq (scRNA-seq) data from CRISPR regulatory screens. The GLM’s response variable, *Y*, is the observed scRNA-seq counts for a gene in each cell, and the predictor, *X_perturb_*, represents the CRISPR perturbation of a putative enhancer for the gene in the same cells. GLiMMIRS also includes covariates to control for cell cycle and percentage of mitochondrial reads, and an offset term to control for the total number of unique molecular identifiers (UMIs) observed in each cell.

So that we could evaluate the performance of GLiMMIRS-base and GLiMMIRS-int, we next developed the GLiMMIRS-sim simulation framework (Fig. 1C). GLiMMIRS-sim accepts user-defined experimental parameters such as number of cells, number of genes, number of distinct regulatory target regions, and gRNA library size. Using these parameters, it then randomly assigns regulatory target regions to genes and gRNAs to cells and also generates reference coefficient values. Reference coefficient values include enhancer effect sizes corresponding to each target region/gene pair and interactions between target regions, which can be used to benchmark GLiMMIRS-base and GLiMMIRS-int.

We used GLiMMIRS-sim to generate a single-cell CRISPRi screen dataset resembling the Gasperini et al.^13^ experimental dataset. We then fit GLiMMIRS-base to the simulated count data so that we could compare the estimated model coefficients to the simulated “reference” values. We compared the estimated coefficients from GLiMMIRS-base to the reference simulation values, using two different settings for the CRISPR perturbation predictor, *X_perturb_*: a ’perturbation probability’ and ’perturbation indicator’. We examined both predictor types because most enhancers in this dataset were targeted by two different gRNAs and the estimated effects of enhancer perturbations can be biased by the presence of low-efficiency guides (Fig. S1, Table S1). In the perturbation probability setting, we defined the value of *X_perturb_* as a function of gRNA efficiency, which is the estimated probability that its intended target is actually repressed in a cell containing the gRNA. In the perturbation indicator setting, we simply set *X_perturb_* to 1 for cells containing a gRNA targeting the enhancer and 0 for all other cells. The perturbation indicator ignores guide efficiency, but is simpler and is the standard approach that is commonly used in the analysis of CRISPR screens^10,13,20,21^.

Upon applying GLiMMIRS-base to the simulated data, we found that the estimated enhancer effects (*β_enhancer_*) correlated well with the reference enhancer effects when we used a perturbation indicator for *X_perturb_* (Pearson *r* = 0.81) and that this correlation improved when we used a perturbation probability for *X_perturb_* (Pearson *r* = 0.86) (Fig. 1D, Table S2). In addition, using the perturbation indicator underestimated the enhancer effects (Fig. 1D). This suggests that a model that accounts for variable guide efficiency using a measure of perturbation probability can obtain more accurate estimates of enhancer activity; however, gRNA efficiency estimates can be noisy, which may impact these estimates^22–25^. To assess this, we performed simulations with varying levels of noise in guide efficiency estimates. In simulations with low to moderate noise, the perturbation probability continued to provide accurate estimates of enhancer effects (Pearson *r* = 0.78 − 0.85). When the efficiency estimate noise was very high, the perturbation probability still yielded reasonable, albeit less accurate, estimates of enhancer effects (Pearson *r* = 0.37) (Fig. S2, Table S3). In summary, GLiMMIRS-base provides accurate estimates of enhancer activity when applied to simulated data and accounting for guide efficiency can improve accuracy when guide efficiency estimates have low or moderate noise.

We then applied GLiMMIRS-base to the Gasperini et al.^13^ dataset and compared the p-values obtained from our model to those from their published analysis, which utilized Monocle 2^26^. We detected a similar number of significant enhancer-gene pairs (560 validated by GLiMMIRS-base out of the 609 reported as significant by Gasperini et al.^13^), but with lower p-values for most of the highly significant pairs. Our p-values are well-calibrated, and when applied to permuted data, where gRNA identities are assigned to different cells, or to shuffled genes, where the candidate enhancers are not near the target gene, the p-value distributions match the null expectation (Fig. 1E, Table S4). These results establish that GLiMMIRS-base provides similar results to the published analysis by Gasperini et al. and may boost power by including additional covariates such as cell cycle scores. Having established the validity of our approach for the simpler scenario of single enhancers acting on single genes, we proceeded to study the regulatory effects of enhancer pairs.

### GLiMMIRS-int detects interactions between pairs of enhancers

To estimate the effects of pairs of enhancers on gene expression, we developed GLiMMIRS-int (Fig. 1B). In this model, we replaced the single enhancer term (*β_enhancer_X_perturb_*) in GLiMMIRS-base with three new terms to represent: 1) the first enhancer in the pair (*β_A_X_A_*); 2) the second enhancer in the pair (*β_B_X_B_*); and 3) an epistatic-like interaction between the enhancers (*β_AB_X_AB_*). We set the values of the *X*_*A*_ and *X*_*B*_ predictors to be perturbation probabilities for the respective enhancers. Likewise, we set the value of *X*_*AB*_to be the probability that both enhancers are simultaneously perturbed (*X*_*AB*_ = *X*_*A*_*X*_*B*_).

To identify pairs of enhancers where interactions could be evaluated by GLiMMIRS-int, we identified pairs of putative enhancers that were targeted for CRISPRi perturbation in the Gasperini et al. experiment where both members of the pair were located within 1MB of a common target gene. We found a total of 795,616 such pairs. Since some cells must contain perturbations to both enhancers (“joint perturbations”) to measure interaction effects, we evaluated the number of cells containing gRNAs targeting both enhancers in these pairs. The majority of enhancer pairs were jointly perturbed in a small number of cells, which limits power to detect interactions; however, we found that 46,166 were jointly perturbed in at least 10 cells (Fig. 2A). We considered this latter set to be “testable” enhancer pairs and restricted our downstream analysis with GLiMMIRS-int to these pairs.

**Figure 2.**
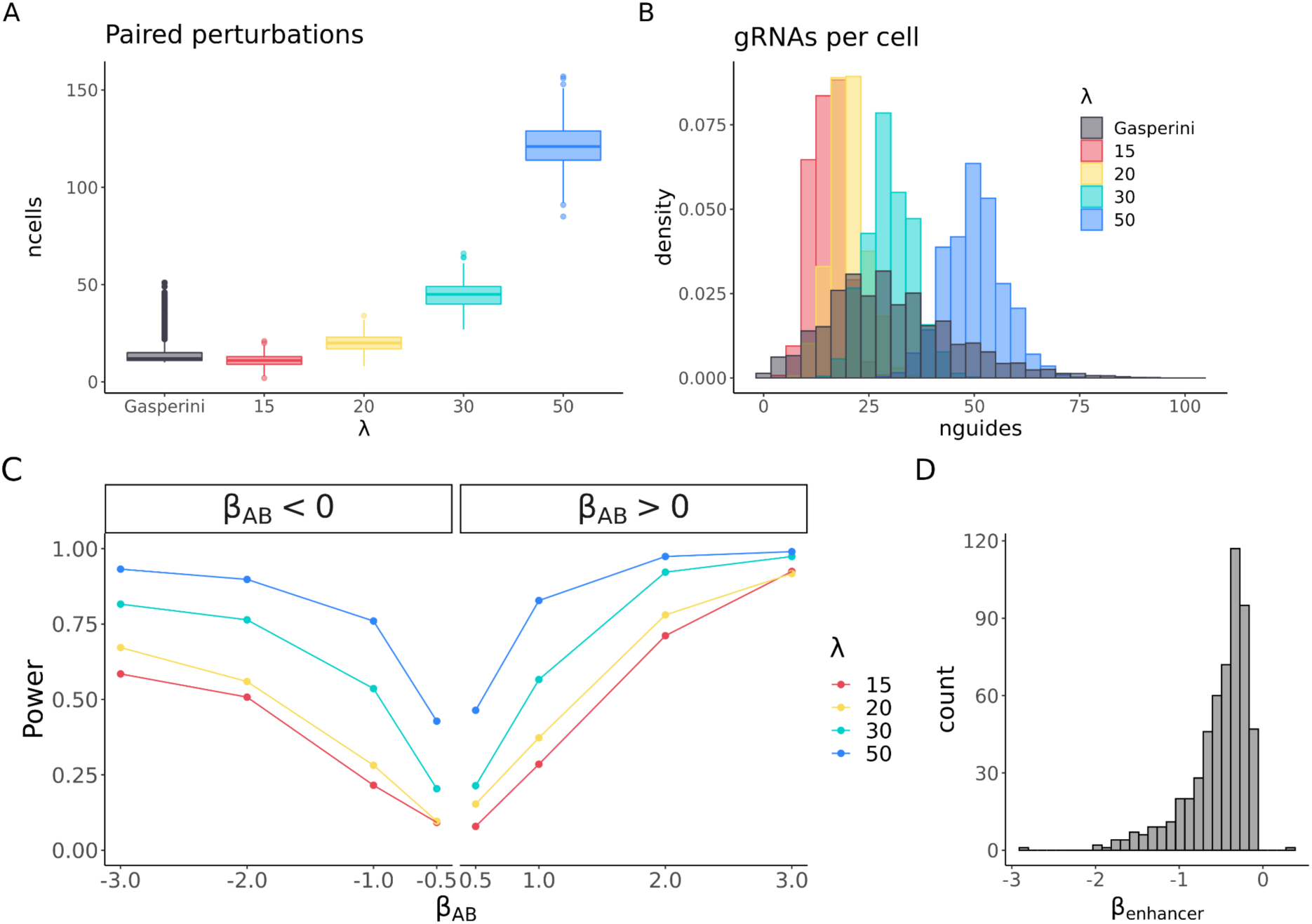
GLiMMIRS-int detects interactions between pairs of enhancers in simulated data. **A)** Boxplots showing the number of cells containing gRNAs targeting both enhancers belonging to a testable pair in the Gasperini dataset (grey) or in the simulated data (colored). To consider an enhancer pair to be “testable”, we required both enhancers to be located within 1MB of a common target gene and for the enhancers to be jointly perturbed in at least 10 cells. **B)** Histogram of the number of unique gRNAs per cell for data simulated with different values of λ (colored) and the Gasperini dataset (grey). **C)** Power to detect interaction effects (y-axis; TPR=true positive rate) in simulated datasets with varying multiplicities of infection (λ) and effect sizes (x-axis) using GLiMMIRS-int. See also Table S5. **D)** Histogram of effect sizes estimated by GLiMMIRS-base for significant individual enhancers in the Gasperini dataset.

To assess our power to detect enhancer interactions with GLiMMIRS-int, we used GLiMMIRS-sim to simulate data over a range of MOIs (λ) and interaction effect sizes (Fig. 2A-B, Table S5). In this simulated dataset, we defined “interacting” enhancer pairs as having an interaction effect on their target gene and “non-interacting” pairs as having individual effects on the target gene but no additional interaction effect. We focused on simulations with MOIs of 15 and 20 since the properties of these simulated datasets most closely resembled the Gasperini dataset (Fig. 2A-B) and restricted our analysis to the testable enhancer pairs that were jointly perturbed in at least 10 cells. As expected, our power to detect interaction effects scaled with the interaction effect sizes and MOI, which increases the number of cells with joint perturbations of both enhancers (Fig. 2A). We found that at MOIs of λ = 15,20, there is reasonable power (7-37%) to detect interactions with modest effect sizes (*β*_*AB*_) of -1.0, -0.5, +0.5 and 1.0 (Fig. 2C, Table S5) and good power (50-78%) to detect strong interactions with effect sizes of -2 or +2. Negative interaction effect sizes as large as -2 are plausible as they resemble the effect sizes we estimated for individual enhancers in the Gasperini et al. dataset (Fig. 2D) and strong interactions might be expected if pairs of enhancers act in a highly redundant or synergistic manner. While our power to detect interactions with these effect sizes is incomplete, we nonetheless expect to detect a substantial number of interactions if they are a common feature of enhancer pairs.

### Enhancers act multiplicatively to control gene expression, but analysis of CRISPR perturbations provide limited evidence for interactions

Next, we applied GLiMMIRS-int to the Gasperini dataset to study enhancer-enhancer interactions. In addition to the testable enhancer pairs defined above, which contained 46,166 pairs and corresponding target genes (of which 5,895 were unique, Fig. S3A), we defined a set of “high-confidence” pairs where each of the individual enhancers showed prior evidence of enhancer activity from the Gasperini study. This high-confidence set contained 264 testable enhancer pairs and corresponding target genes (of which 94 were unique, Fig. S3B), where each of the individual enhancers had a previously reported effect on the target gene based on the Gasperini analysis.

We first examined whether the combined effects of multiple enhancers on gene expression were better described by a multiplicative or additive model. To this end, we fit two versions of GLiMMIRS-int to the 264 enhancer pairs and their target genes in the high-confidence set: an additive model, in which we used an identity link function; and a multiplicative model, in which we used a log link function. We then compared the model fits with Akaike Information Criterion (AIC), an approach similar to that used by Dukler et al.^5^ to compare different enhancer activity models. In all cases, the multiplicative model provided a better fit to the data, indicating that the combined effect of enhancers is better described by a multiplicative model (Fig. 3A). Thus, we used the multiplicative form of GLiMMIRS-int in all subsequent analyses.

**Figure 3.**
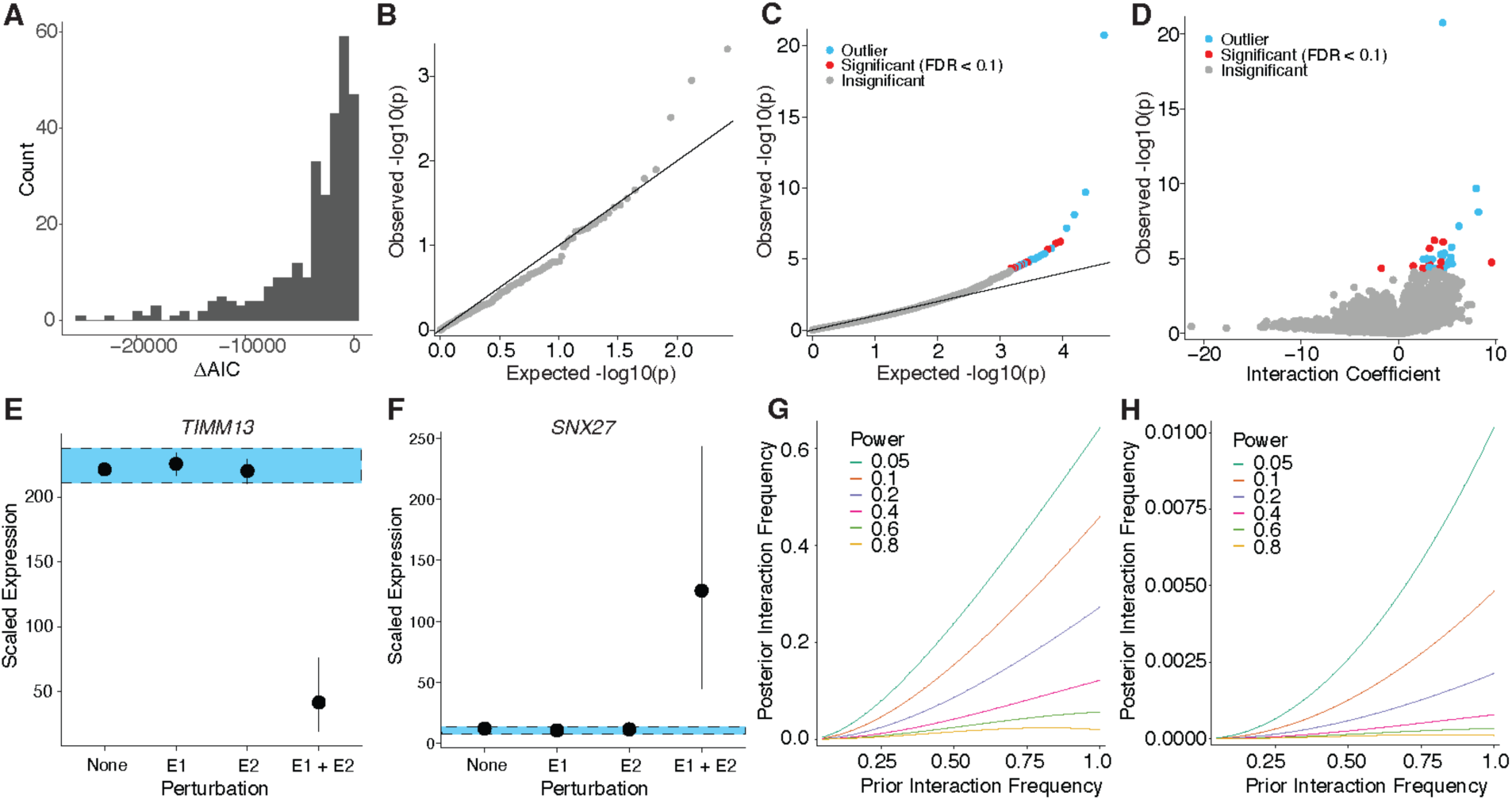
Enhancers act multiplicatively to control gene expression, but analysis of CRISPR perturbations provide limited evidence for interactions. **A)** Distribution of *ΔAIC*, the difference in Akaike Information Criterion between the best fitting model and the lesser model for 264 high confidence enhancer pairs and corresponding target genes from Gasperini et al. In every case evaluated, the multiplicative model fit better than the additive model. **B)** Quantile-quantile (QQ) plot of interaction coefficient p-values for 264 high confidence enhancer pairs, where each individual enhancer had significant effects on the target gene expression. None of the interaction coefficients were significant (Benjamini Hochberg FDR < 0.1). **C)** QQplot of 46,166 enhancer pairs in the entire testable set, where each constituent enhancer did not necessarily have a significant effect on gene expression. Significant interaction coefficients (FDR<0.1) are blue if one of the jointly perturbed cells was an outlier by Cook’s distance and red otherwise. **D)** Volcano plot of interaction coefficients for the 46,166 enhancer pairs in the entire testable set. **E)** An enhancer pair with a significant negative interaction on the expression of *TIMM3*. The Y axis is *TIMM13* expression estimated in cells lacking perturbations to either enhancer (None), cells with perturbations of one enhancer (E1, E2) and cells with joint perturbations of both enhancers (E1+E2). The blue rectangle is the expected expression in joint perturbation condition under the null model of multiplicative enhancer effects (90% CI estimated from 100 bootstraps). Whiskers are 90% CIs of expression estimates (from 100 bootstraps). **F)** A gene (*SNX27)* and enhancer pair with a significant positive interaction. **G)** Posterior estimate of frequency of enhancer pairs with interactions estimated from the dataset of 264 high-confidence enhancer pairs. Each line corresponds to a different power to detect interactions. X-axis is the prior belief in enhancer interaction frequency. **H)** Posterior estimate of frequency of enhancer pairs with interactions estimated from the 46,166 enhancer pairs in the entire testable set.

We applied GLiMMIRS-int to the 264 enhancer pairs in the high-confidence set and observed no significant interaction terms (Likelihood Ratio Test, FDR<0.1) (Fig. 3B, Table S6). We then applied GLiMMIRS-int to the 46,166 enhancer pairs in the entire testable set and identified 31 significant interaction term effects (Likelihood Ratio Test, FDR<0.1) (Fig. 3C, Table S7). Of these significant interaction terms, 30 out of 31 were positive (Fig. 3D, Table S7). To better understand these significant interactions, we examined the distribution of single-cell RNA-seq UMI counts for the target genes, focusing on the cells that received guides targeting both of the corresponding enhancers. In the majority of cases, we observed a small number of outlier cells with very high UMI counts (Fig. S4A). Since GLM coefficients and p-value estimates can be sensitive to outliers, we computed Cook’s distance for each cell containing a joint perturbation. Cook’s distance quantifies the influence of a single observation on the coefficient estimate. The outlier cells had large Cook’s distances, indicating that they disproportionately affected coefficient estimates for the interaction effect, *β*_*AB*_(Fig. S4B). Therefore, we removed enhancer pairs where any of the cells containing a joint perturbation had an extreme Cook’s distance (max / mean ratio>5). After applying this filter, 11 out of 31 significant signals remained (Table S8).

The remaining interactions included one negative enhancer interaction for the gene *TIMM13* (Fig. 3E). In this case, a reduction in expression was observed only when both of the enhancers were jointly perturbed, potentially representing an example of “enhancer redundancy”^6,27^. However, we believe that this enhancer pair is unlikely to be a true case of enhancer redundancy as neither of the targeted candidate enhancers are marked by the canonical enhancer histone modification H3K27ac in K562 cells (Fig. S5), and they are located very far from one another (>988 kb) in different topological associating domains (Fig. S6).

The other 10 remaining significant interactions were all positive and generally followed a pattern in which the expression of the target gene was low in the absence of any enhancer perturbations and elevated in jointly perturbed cells, as if the gene had become de-repressed (Fig. 3F). De-repression of gene expression is not an expected response to targeting of enhancers by CRISPRi and could potentially be due to regulatory effects on other genes that exert downstream effects on the target genes.

Based on our analysis, evidence for interactions between enhancers in the Gasperini dataset is very limited. In the cases where we do identify significant interactions, they do not have characteristics that would be expected given previously postulated “synergistic” or “redundant” enhancers^6,27^. To quantify the possible frequency of enhancer interactions throughout the genome, we estimated the posterior frequency of enhancer interactions given a range of detection probabilities corresponding to our power analysis (Fig. 2C) and given different priors for the frequency of enhancer interactions (Fig. 3G-H). For example, with a prior enhancer interaction frequency of 25% and a detection probability of 20%, which roughly corresponds to our power to detect moderate interactions (effect size = -1), the posterior enhancer interaction frequency for the high-confidence set of enhancers is 2.4% (Fig. 3G). Even if we assume a very high prior enhancer interaction frequency of 50%, our posterior estimate is only 8.6%. Since a detection probability of 20% corresponds roughly to our power to identify moderate interactions (effect size -1.0), we infer that the frequency of enhancer interactions of moderate strength is likely less than 10%. If we consider a low detection probability of 5% corresponding to weak interactions, the posterior interaction frequency estimates depend more strongly on the priors and are 7.9% for a prior of 25% and 24% for a prior of 50% (Fig. 3G). Thus, we have less certainty in the frequency of weak enhancer interactions but conclude that they are not likely to be extremely common.

For our entire set of 46,166 testable enhancer pairs, the posterior estimates of enhancer interactions are far lower (Fig. 3H). Here, the frequency of interactions is expected to be low because most of the individual enhancers included in this analysis do not have evidence for independent effects on the expression of their corresponding target genes. Nonetheless, this analysis implies that cases of enhancer redundancy, where enhancers’ effects on expression are only observed when two enhancers are jointly perturbed, must be uncommon.

### Validation with independent CRISPR perturbation dataset and in-silico perturbation experiments

To examine enhancer interactions in an independent dataset, we next applied GLiMMIRS to a CRISPRi regulatory screen performed by Morris et al^28^. This study targeted candidate regulatory sequences in K562 cells that were implicated in genome-wide association studies of blood cell traits. We focused our analysis on the *PTPRC* locus because 6 out of the 9 candidate enhancers targeted near this gene were reported to affect its expression. We re-analyzed this dataset using GLiMMIRS-base and confirmed that those same 6 enhancers had significant effects on *PTPRC* expression (Table S9; Bonferroni-corrected p < 0.1). We then tested all 36 possible combinations of pairs from the 9 candidate enhancers for interactions using GLiMMIRS-int and found two enhancer pairs with significant interaction terms (Bonferroni-corrected p < 0.1). However, our Cook’s distance analysis indicated that both significant interactions were driven by outliers (Fig. 4A, Table S10). Thus, we conclude that there is no evidence for enhancer interactions among the 9 candidate enhancers for *PTPRC*.

**Figure 4.**
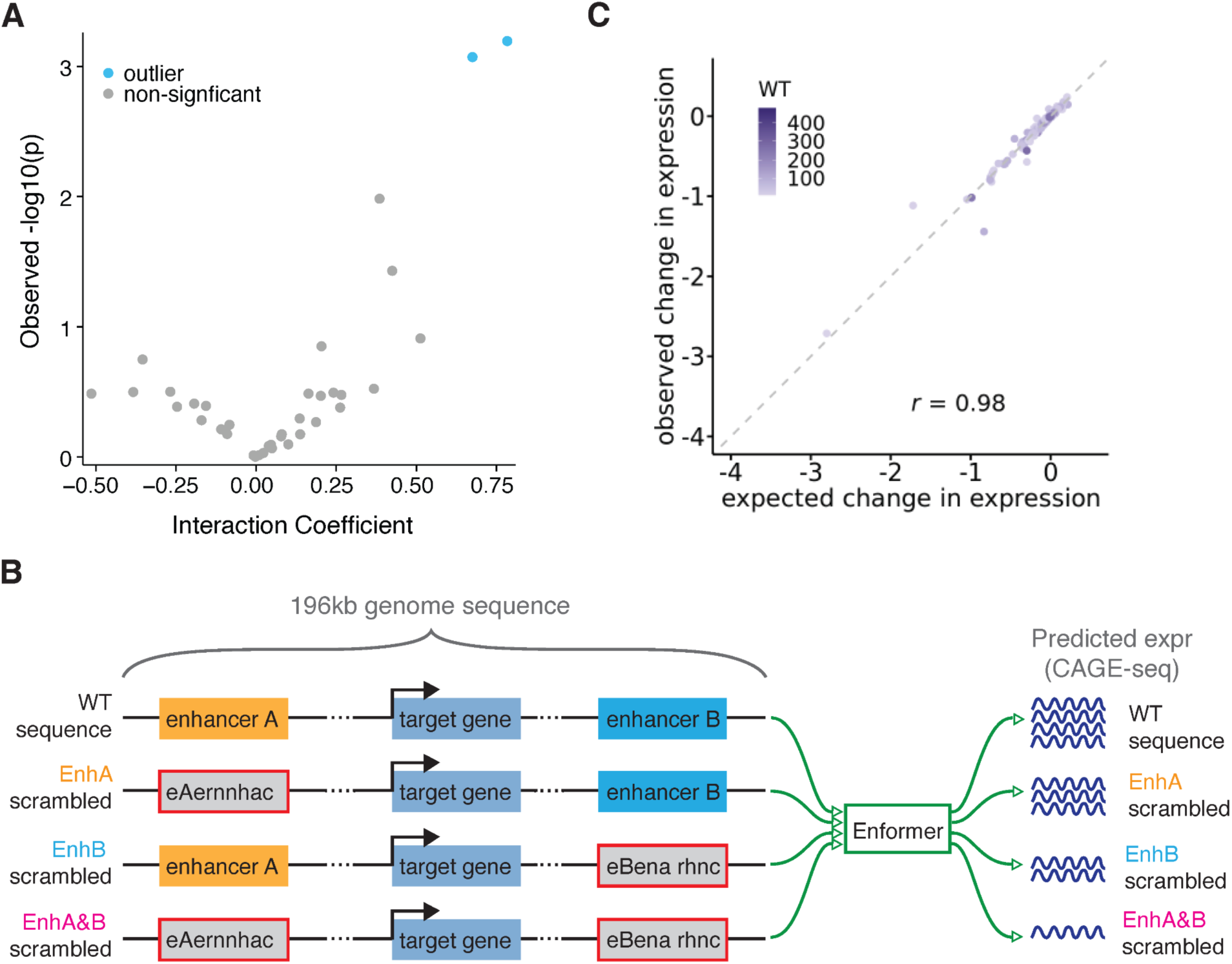
No evidence for enhancer interactions from an independent CRISPRi dataset and in-silico perturbations. **A)** Volcano plot of interaction coefficients and -log10(p values) estimated by GLiMMIRS-int from the Morris et al. CRISPRi perturbation dataset. All 36 possible pairs of enhancers for the 9 candidate enhancers for *PTPRC* were tested. Both significant interactions were driven by jointly perturbed cells with outlier expression levels. **B)** Schematic of in silico perturbation strategy. We input 196kb sequences into Enformer, where each input sequence contained both candidate enhancers and the target gene. We generated predictions from unperturbed wild type (WT) sequences, sequences with the first enhancer (EnhA) scrambled, sequences with the second enhancer (EnhB) scrambled, and sequences with both enhancers scrambled (EnhA&EnhB). **C)** We compared the Enformer-predicted change in expression of the double mutant (EnhA&EnhB) to the expected expression under a multiplicative model with no enhancer interactions. We analyzed 1372 enhancer pairs and genes where Enformer-predicted WT expression >10 and quantified the change in expression as the log ratio of mutant expression to WT expression. The expected expression for the double mutant was computed from the Enformer-predicted expression of the single (EnhA and EnhB) mutants under a multiplicative model of activity. The shading of points corresponds to the Enformer-predicted WT expression level.

As an orthogonal approach to examine interaction effects between enhancer pairs, we used Enformer^29^ to perform in-silico perturbation experiments. Enformer is a state-of-the-art deep learning model that can accurately predict gene expression from long genomic sequences (∼200kb). To use Enformer to estimate the effects of enhancers on gene expression, we provided ’wild type’ sequences and synthetic sequences where regions corresponding to enhancer pairs from the Gasperini dataset ^30^ were shuffled (Fig. 4B). We then compared Enformer’s predictions when individual enhancers were shuffled versus when both enhancers were shuffled simultaneously (Fig. 4B, Table S11). The Enformer-predicted changes in gene expression when both enhancers in a pair were simultaneously perturbed were strongly correlated with the expected change in gene expression under a multiplicative model of activity given the predictions for individual enhancer perturbations (Pearson *r* = 0.98; Fig. 4C). This strong correlation is consistent with a model of activity that does not require interaction effects between enhancers and provides further evidence that interactions between pairs of enhancers are uncommon.

## Discussion

CRISPR perturbations provide a new way to measure how combinations of enhancers regulate gene expression. We reanalyzed data from a single-cell CRISPRi experiment designed to map enhancers to the genes that they regulate. Since this dataset transduced gRNAs with a high MOI, multiple enhancers within 1MB of the same gene were sometimes perturbed within the same cells, making it possible to analyze the joint effects of multiple enhancers on gene expression. Our analysis of two CRISPRi datasets (Fig. 3,4A) and additional in silico sequence perturbations (Fig. 4B,C) support a model in which enhancers act multiplicatively to control gene expression. Such a model was previously proposed by Dukler et al.^5^, whose analysis of enhancers supported either a logistic or multiplicative model of regulatory activity over an additive model; however, this study was limited to examining enhancers at only two loci. Likewise, a subsequent study proposed the now well-known ABC model of activity based on a perturbation experiment targeting candidate enhancers for 30 genes; however, this study perturbed each enhancer individually and so its conclusions about their regulatory activity cannot be extended to the phenomenon of joint enhancer effects^32^.

A novel aspect of our study is that we analyzed joint perturbations of thousands of pairs of enhancers. Under a multiplicative model of enhancer activity, we analyzed pairs of enhancers that were jointly perturbed by CRISPRi and found that the experimental results resemble those expected under the null hypothesis of no enhancer interactions (Fig. 3B,C; Fig. 4A). A limitation of the Gasperini dataset that we analyzed is that even with a high MOI and a large number of sequenced cells, only a subset of enhancer pairs could be interrogated. Specifically, we tested 46,166 of a possible 795,616 enhancer pairs in the Gasperini dataset because most were not simultaneously perturbed in a sufficient number of cells. Furthermore, our power to detect weaker interactions was limited. For example, we only had 21.5% power to detect interactions with an effect size of -1.0 under a simulated MOI of λ = 15 (Table S5). Many of these power limitations could be overcome through CRISPRi experiments designed specifically to probe enhancer interactions. For example, a high MOI CRISPRi experiment could be performed in which a much smaller number of candidate enhancers are targeted so that testable pairs are frequently perturbed simultaneously in the same cells. Multiple guides could also be transduced on the same vectors so that nearby enhancers are guaranteed to be targeted in many cells^31^. This type of approach was recently used to estimate enhancer interactions at the MYC locus^9^. Another advantage of providing multiple guides on the same vector is that enhancer interactions could be interrogated with low MOI experiments, which would circumvent two potential issues with high MOI experiments. First, Cas9 competition for gRNAs may reduce experimentally observed enhancer effect sizes, and second, high MOIs may induce unintended cellular responses that alter gene expression.

Further limitations of our analysis are that both of the CRISPRi datasets that we analyzed were from K562 cells and it is possible that enhancer interactions are more prevalent under dynamic conditions or in different cell types. Interactions may also be more common among enhancers that are clustered into ’super-enhancers’ which were not specifically interrogated by the Gasperini dataset.

Despite the above limitations, our results argue against the presence of strong epistatic interactions between enhancers. If such interactions do exist, they must be infrequent, of small effect, or restricted to specific cell types or conditions. How can these observations be reconciled with prior reports of enhancer redundancy or synergy? A possible explanation is that if an additive model is assumed, then interaction terms are often required because the combined effects of multiple enhancers are greater (synergistic) or less than (redundant) than expected under an additive model. However, these deviations from additivity may be naturally accounted for by a multiplicative model without the need for an interaction term. For example, under a multiplicative model, perturbation of a weak enhancer may have a small or negligible effect on expression on its own but a much more substantial effect when combined with a perturbation to a strong enhancer. An additive model would require an interaction term to describe these results and the enhancers would appear to be ’redundant’.

A recent study by Lin et al. analyzed enhancer interactions at the *MYC* locus using pairs of CRISPR guides and reported additive interactions between nearby enhancers and synergistic interactions between distant enhancers^9^. Our results are difficult to compare with those from Lin et al. for two reasons. First, the high-throughput screen in Lin et al. was performed using cell proliferation rather than gene expression as readout, thereby assuming that proliferation was proportional to *MYC* expression. Second, while Lin et al. examined how selected pairs of enhancers affect the expression of *MYC* and other genes, their analysis relied on log relative expression obtained by RT-qPCR, which may not be directly comparable to expression estimated from scRNA-seq UMI counts.

Future studies examining enhancer interactions will benefit from GLiMMIRS, which uses a generalized linear model that accounts for guide efficiency, differences in per-cell sequencing depth and several covariates. We note that it is important to consider a multiplicative model as the baseline expectation when looking for enhancer interactions, and when interactions are identified it is important to consider the possibility that the results are driven by a small number of outlier cells. To increase power to detect weak interactions, CRISPR experiments that are specifically designed to examine enhancer interactions are desirable. Our study motivates the further study of enhancer interactions in more cell types and conditions, to which GLiMMIRS can be applied to yield novel insights into regulatory element interactions and their effects on transcription.

## STAR★Methods

### CRISPRi perturbation of NMU enhancers

We identified two target sites of interest, A and B, for the gene *NMU*, each of which was targeted by two gRNAs in the Gasperini et al. experiment (A1 and A2 targeting enhancer A; B1 and B2 targeting enhancer B). Pairs of gRNAs were designed by FlashFry to target enhancers A and B at the same time, using 2 gRNAs per site. The gRNA pairs included the following: NMU_tss+NMU_tss (positive control), Safe_harbor (SH)+SH (negative control), A_sgRNA1+SH, A_sgRNA2+SH, SH+B_sgRNA1, SH+B_sgRNA2, A_sgRNA1+B_sgRNA1, A_sgRNA1+B_sgRNA2, A_sgRNA2+B_sgRNA1, A_sgRNA2+B_sgRNA2. Pairs of gRNAs were cloned into pLV-dCas9-KRAB-puro (Addgene #71236) following published methods^33,34^. Briefly, DNA oligos carrying pairs of guides were synthesized by IDT and cloned into pLV-dCas9-KRAB-puro plasmids by Gibson assembly reactions. Lentivirus was generated by co-transfecting the plasmid with PsPAX2 (Addgene #12260) and pMD2.G (Addgene #12259) in 293FT cells obtained from the Salk Institute Stem Cell Core. Lentivirus was harvested 48h post transfection. K562 cells (ATCC #CCL-243) were transduced by the lentiviruses using spinoculation. 72h after transduction, K562 cells with viral genome integration were selected by puromycin for 48h. Total RNA from live K562 cells was extracted and reverse transcribed using SuperScript IV First-Strand Synthesis System (Thermo Fisher Scientific #18091050) with random hexamers. *NMU* expression was quantified by reverse transcription quantitative PCR (RT-qPCR). CRISPR gRNA designs and PCR primers used in experiment can be found in Table S1.

### Gasperini et al. Dataset

Data from the at-scale screen in the Gasperini et al. study are available at GEO accession number GSE120861. We downloaded guide spacer sequences from Supplementary Table 2 of their paper^13^. The single-cell RNA-seq expression matrix from the at-scale screen was downloaded from the GEO file ‘GSE120861_at_scale_screen.exprs.mtx.gz’. The cell barcodes were determined from the GEO file ‘GSE120861_at_scale_screen.cells.txt.gz’. Gene names were determined from the GEO file ‘GSE120861_at_scale_screen.genes.txt.gz’. Covariate information as well as cell-guide mapping information was determined from the GEO file: ‘GSE120861_at_scale_screen.phenoData.txt.gz’. Differential expression results were downloaded from ‘GSE120861_all_deg_results.at_scale.txt.gz‘ to determine candidate enhancer pairs from the 664 enhancer-gene links. Enhancer-guide assignments from the at-scale screen were downloaded from ‘GSE120861_grna_groups.at_scale.txt.gz‘ to find candidate enhancer pairs for the larger low-confidence set.

### Computing guide efficiencies

We first collected the 13,189 guide RNA sequences used in the at-scale screen previously published by Gasperini et al.^13^, which were published in Supplementary Table 2 of their study. We then appended ‘NGG’ to each spacer sequence for compatibility with GuideScan 2.0^35^. We then used the GuideScan 2.0 gRNA sequence search tool (https://guidescan.com/grna) with the organism ‘hg38’ and the enzyme ‘cas9’ parameters to predict efficiencies for the guide RNA spacer sequences. We used the “Cutting.Efficiency” values outputted from GuideScan as our guide efficiency values. These values are equivalent to the “Rule Set 2” scores defined by Doench et al. 2016 and can be used to predict the on-target activity of gRNAs in CRISPRi screens^23^.

Out of the 13,189 guide RNA sequences, 762 guide RNAs were designed to target transcription start sites, 101 guide RNAs were designed as non-targeting controls, 14 guide RNAs were designed as positive controls targeting the globin locus, and the remaining 12,312 guide RNAs were designed to target candidate enhancer sequences.

From the 12,312 enhancer-targeting guide RNAs, 2,331 guides did not have a guide efficiency value because GuideScan does not compute scores for guides for which it cannot find a match or for which there are multiple matches within edit distance 1 (Table S12). We excluded these 2,331 guide RNA sequences from downstream analysis so that in total 9,981 guides were used for our downstream analysis.

### Computing cell cycle scores for Gasperini et al

Cell cycle scores were computed from the single-cell RNA-sequencing gene expression matrix from the at-scale screen previously published by Gasperini et al.^13^ using the Seurat^36–40^ R package.

Since the Seurat R package uses gene names from the HUGO Gene Nomenclature Committee, gene names were converted from their Ensembl Gene ID to HGNC symbol (https://www.genenames.org/) using the biomaRt^41^ tool from Ensembl^42^ with the “hsapiens_gene_ensembl” dataset. Of the 13,135 genes in the at-scale expression matrix, 351 genes were not recognized by BioMart and 571 genes did not successfully map from Ensembl Gene ID to HGNC symbol. For the total 922 genes that could not be mapped from Ensembl Gene ID to HGNC symbol, the Ensembl Gene ID was imputed as the gene name for downstream analysis with Seurat (Table S13).

To determine cell cycle scores, we used pre-defined sets of genes associated with S and G2M phases from the Seurat library. We log-normalized the data, identified variable features, and scaled the expression matrix using functions defined in Seurat. We then used the cell cycle scoring function with the predefined S and G2M gene sets in Seurat to compute cell cycle scores for each cell in the at-scale screen. All Seurat functions were run with default parameters (Tables S14, S15).

### Model fitting and implementation

All models were fitted by maximum likelihood using the ‘glm.nb()’ function from the MASS package in R^43^. Both GLiMMIRS-base and GLiMMIRS-int use a negative binomial generalized linear model with a log link function. The additive model was implemented with an identity link function.

### Defining a baseline model for a single enhancer acting on a single target gene

Our baseline model tests for the simple case where a single enhancer acts on a single gene. The model is a generalized linear model which assumes a log link function and that the single-cell RNA-seq UMI counts of each gene follow a negative binomial distribution. In other words, *y*∼*NB*(*µ*, *ϕ*), where *y* represents the scRNA-seq UMI counts of the genes, *ϕ* represents the dispersion parameter of the negative binomial distribution, and *µ* is the mean parameter of the negative binomial distribution. The mean parameter is specified by a linear predictor passed through an exponential (inverse log-link) function: *µ* = *exp* (*β*_0_ + **β*_enhancer_X_perturb_* + *β*_*S*_*X*_*S*_ + ^*β*^_*G*2*M*_^*X*^_*G*2*M*_ ^+ *β*^_*m*i*to*_^*X*^_*m*i*to*_ ^+ *β*^_g*RNAs*_^*X*^_g*RNAs*_ + *β*_*batc*h_^*X*^_*batc*h_ ^+ *ln*(*s*)^).

In this expression, we have gene-specific coefficients and cell-specific predictor values. *β*_-_is the intercept and represents the baseline gene expression before the influence of any other relevant factors on gene expression. **β*_enhancer_* represents the effect of a perturbed target site (putative enhancer) on its target gene. *β*_*S*_ and *β*_*G*2*M*_ are coefficients that represent the effect of the S and G2M cell cycle states, respectively. *β*_*m*i*to*_ is a coefficient representing the effect of percentage of mitochondrial reads. Finally, *β*_g*RNAs*_ is a coefficient representing the effect of the total number of gRNAs observed within a given cell. *β*_*batc*h_ is a coefficient representing the effect of the prep batch, from the Gasperini et al. 2019 experiment. We incorporate measures of guide efficiency in the variable *X_perturb_*. This variable is calculated for each cell based on the efficiencies of every gRNA targeting the target site being modeled which are present in the cell. Specifically, *X_perturb_* is calculated for any given cell and target site as 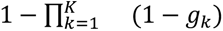, where *K* is the total number of gRNAs targeting the target site found in the cell and *g*_*k*_ is the efficiency of the *k*^*t*h^ gRNA. Because we interpret guide efficiency as the probability that a gRNA successfully perturbs its designated target site, the expression for *X_perturb_* can be interpreted as the joint probability of a perturbation in a given cell based on all of the gRNAs targeting the site that are present in that cell. *X*_*S*_ and *X*_*G*2*M*_ are S and G2M cell cycle scores, respectively, for each cell. *X*_*m*i*to*_ is the percentage of mitochondrial reads in a cell. *X*_g*RNAs*_ is the total number of gRNAs observed in a cell. *X*_*batc*h_ is the prep batch (from Gasperini et al. 2019). Finally, *s* is an offset term for the model that serves as a scaling factor controlling for variable sequencing depth across cells. It is calculated as 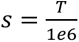, where *T* is the total scRNA-seq UMIs in a cell summed across all genes in the expression count matrix. Prior to fitting the models, we added a pseudocount of 0.01 to the scRNA-seq counts of the gene being modeled for all cells to prevent inflation of coefficients (see section: Defining a model for an enhancer pair acting on a single target gene).

### Simulating data for single enhancers acting on single genes

To begin, we define some simulation parameters, including the total number of cells, *C*; the total number of genes, *G*; the total number of target sites, *N*; and the number of gRNAs targeting each site, *d*. Note that the total number of target sites, *N*, is also the total number of target genes, as this simulation assumes that each target site is a unique enhancer for a unique gene. To generate a simulated dataset, we need to simulate sets of coefficient values for each gene (*β*_-_, **β*_enhancer_*, *β*_*S*_, *β*_*G*2*M*_, *β*_*m*i*to*_) as well as corresponding variable values for each cell (*X_perturb_*, *X*_*S*_, *X*_*G*2*M*_, *X*_*m*i*to*_, and scaling factor *s*). We also need to simulate the gRNA library and assign gRNAs to cells, as well as assign guide efficiencies to gRNAs (which will be used to calculate *X_perturb_*). These values are used to calculate a value of *µ* for defining a negative binomial distribution from which simulated counts for a given gene will be drawn. Specifically, *µ* = *exp*(*β*_-_ + **β*_enhancer_X_perturb_* + *β*_*S*_*X*_*S*_ + *β*_*G*2*M*_*X*_*G*2*M*_ + *β*_*m*i*to*_*X*_*m*i*to*_ + *ln*(*s*)). The terms for total gRNA counts per cell and batch are omitted from the simulation for simplicity, and are also omitted when fitting the baseline model to the simulated data. The dispersion parameter for the negative binomial distribution will be constant across all genes and estimated from the empirical data. For the simulated dataset described in our paper, we used values of *G* = 13000, *N* = 1000, *d* = 2.

We first simulated values of **β**_0_ for each gene. To do this, we randomly selected a subset of 1,000 genes and 10,000 cells from the Gasperini et al. 2019 at scale experiment and fit the counts for these genes to negative binomial distributions using maximum likelihood estimation (MLE). Specifically, we define the mean parameter of the negative binomial here as *µ* = *exp*(*β*_0_ + *ln*(*s*)). Note that here *s* is calculated from the total counts for the gene across the subset of 10,000 cells using the formula defined in the previous section. This simplified model has no covariates, but does account for the scaling factor, as the goal is to simply get a sense of what coefficient values reflect the empirical data. After modeling the counts from the random subset of data, we visualized the distribution of estimated *β*_0_-(from which *µ* is calculated) and dispersion parameters for each gene tested. From what we observed, we picked a fixed dispersion value of *ϕ* = 1.5 for defining the negative binomial distribution for generating simulated count data. We also observed that the distribution of *β*_0_-estimated from the subset of the at scale experiment were roughly normally distributed. Therefore, we fit these estimated *β*_0_-values to a normal distribution with MLE to obtain parameters for defining a normal distribution from which to sample *β*_-_ values for the simulated dataset. We obtained parameters for the normal distribution of *µ* ≈ 2.24 and *σ* ≈ 1.8, so we sampled *G* times from *N*(*µ* = 2.24, *σ* = 1.8) to yield baseline coefficients for all the genes in the simulated dataset.

To assign guides to cells, we first determined the number of gRNAs in each cell in our simulated dataset by sampling from a Poisson distribution defined as *Pois*(λ = 15). This value of λ comes from the fact that in the Gasperini et al. 2019 experiment, they observed a median of approximately 15 unique gRNAs per cell. Thus, we sampled *C* times from the distribution defined by *Pois*(λ = 15) to obtain the number of unique gRNAs in each cell. To assign gRNAs to each cell, we sampled *g* times without replacement from the set of all gRNAs in our library, where *g* is the total number of gRNAs in each cell (determined in the previous step) and the gRNA library is denoted as a sequence of integers 1,2, . . . , *dN*. Information about which gRNAs are found in which cells are stored in a one hot encoded matrix.

We defined guide efficiency for each gRNA by sampling from a left-skewed Beta distribution, to represent the fact that an experimental design would select for gRNAs with higher efficiencies). For our simulation we used a Beta distribution defined as *Beta*(*a* = 6, *b* = 3).

Next, we created a mapping of gRNAs to target genes. For each target site, or putative enhancer, we randomly select an integer from 1,2, . . . , *G* to represent the target gene of the candidate enhancer (indexers are used as gene identifiers). This is done without replacement to simulate a case where we are attempting to study enhancers of distinct genes, and yields a vector of length *N*, which we will replicate *d* times to yield a complete mapping of gRNAs to target genes. In this vector of length *Nd*, the index of a given value in the vector represents the gRNA identifier.

Enhancer effect sizes are represented by the coefficient **β*_enhancer_* and are assigned on a per-gene basis. These values represent the effect that an enhancer has on the expression of its target gene. To do this, we sampled from a gamma distribution and multiplied the values by -1 to yield a negative value, representative of the expectation that successful repression of an enhancer will most likely decrease target gene expression. We wanted the values to be on a comparable scale with the expected baseline expression, *β*_-_, while also not being so small that they would be difficult for the model to detect changes in expression. We chose to sample values of **β*_enhancer_* from a gamma distribution defined by *Γ*(*α* = 6, *σ* = 0.5), and all values drawn from the distribution were multiplied by -1 to represent a negative effect on target gene expression, which is the expectation when an enhancer is repressed.

*X_perturb_* is calculated for each cell as a function of guide efficiencies for the gRNAs targeting the putative enhancer of interest found in that cell. Specifically, it is calculated for each cell as 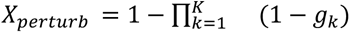 where *K* is the total number of gRNAs targeting the putative enhancer of the gene being simulated/modeled that are present in the cell and *g*_*k*_ is the guide efficiency of the *k*th gRNA in this set of targeting gRNAs. *X_perturb_* = 0 when *K* = 0. We compared the performance of using this variable in our model against the performance of using a binary indicator variable that simply represents the presence of any gRNA targeting the gene being simulated/modeled in each cell.

We generated cell cycle scores for each cell in our simulated dataset using a similar approach to the one we used for sampling *β*_-_values. That is, we first fit models to the empirical data to identify a distribution to draw simulated values from such that they would reflect the distribution of the real data. We first calculated S and G2M cell cycle scores for the empirical data using Seurat’s CellCycleScoring() function^36–40^. We observed that while the S cycle scores calculated from the empirical data appeared to be normally distributed, the G2M scores appeared to show a right skewed distribution. Thus, we fit the empirical S cycle scores to a normal distribution and the empirical G2M scores to a skew normal distribution with MLE. We used the estimated parameters to define distributions for sampling S and G2M scores for the simulated dataset. Specifically, we sampled *C* times from a normal distribution defined by *N*(*µ* = −1.296*e* − 3, *σ* = 0.11) and a skew normal distribution defined by *N*(*ζ* = −0.256, ω = 0.312, *α* = 6.29, *τ* = 0) to obtain simulated S and G2M scores, respectively.

We generated corresponding values of *β*_*S*_ and *β*_*G*2*M*_ by sampling from the same distribution used to generate the enhancer effect sizes, or the gamma distribution defined by *Γ*(*α* = 6, *σ* = 0.5).

Percentage of mitochondrial DNA per cell is simulated using the same approach used to simulate the cell cycle scores and baseline expression values (*β*_-_). We fit to the empirical percentages of mitochondrial DNA per cell. We fit to a beta distribution using MLE, and used the resulting parameter estimates to define a new beta distribution from which we sampled simulated values of percentage of mitochondrial DNA. This beta distribution was defined as *Beta*(*a* = 3.3, *b* = 81.48).

Coefficients for the effect size of percentage of mitochondrial DNA, *β*_*m*i*to*_, were simulated per gene by sampling from the same gamma distribution used to sample the other coefficients (**β*_enhancer_*, *β*_*S*_, *β*_*G*2*M*_). This is the gamma distribution defined as *Γ*(*α* = 6, *σ* = 0.5).

Finally, we simulated scaling factor values, *s*, for each cell in our simulated experiment, which were used to calculate values of *µ* for simulating counts for each gene. To do this, we simulated values of *T*, or total counts per cell, for each cell by sampling from a Poisson distribution defined by *Pois*(λ = 50000), where 50000 is the expected number of reads observed in each cell in a scRNA-seq experiment.

### Simulating noisy guide efficiencies

The noisy guide efficiency estimate, *w*, for a given gRNA in our simulated dataset was sampled from a new Beta distribution parameterized by *a*′ and *b*′, which are calculated from the “true” simulated guide efficiency for the gRNA, *w*, and a dispersion-controlling constant *D*. We wanted the noisy guide efficiency to be sampled from a Beta distribution whose mean is equivalent to the “true” guide efficiency value; thus, 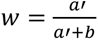. We defined the dispersion-controlling constant *D* as *D* = *a*′ + *b*′. From this, it follows that *a*′ = *Dw* and *b*′ = *D* − *a*′. Like so, we calculated values of *a*′ and *b*′ from which to draw the noisy guide efficiency estimate for a given gRNA in our simulated guide library. The magnitude of *D* is inversely proportional to the amount of noise (Fig. S2A,B).

### Fitting baseline model to simulated data

To fit the baseline model to simulated data, we used a negative binomial GLM with a mean defined by the same log-link function described for generating simulated counts: *µ* = *exp*(*β*_0_ + **β*_enhancer_X_perturb_* + *β*_*S*_*X*_*S*_ + *β*_*G*2*M*_*X*_*G*2*M*_ + *β*_*m*i*to*_*X*_*m*i*to*_ + *ln*(*s*)). Models were fitted by MLE. Each model can be described as *y* = *NB*(*µ*, *ϕ*), where *y* is the simulated counts for the gene being modeled, and all variable values (*X_perturb_*, *X*_*S*_, *X*_*G*2*M*_, *X*_*m*i*to*_) come from the per-cell values from the simulated dataset. We omit *β*_g*RNA*_ when fitting to the simulated data for simplicity.

### Evaluating performance of baseline model on simulated data

Our simulated dataset had *N* target sites, or genes that were regulated by an enhancer perturbed in the experiment. For each of these genes, we computed the Pearson correlation (*r*) between the estimated coefficients, derived from fitting the baseline model to the simulated data, and the “true” (or reference) coefficients, which were generated for the simulation and used to parameterize the distribution from which the simulated counts were drawn. These results are summarized in Table S2 for the continuous vs. indicator forms of *X_perturb_* and in Table S3 for the three different sets of noisy simulated guide efficiencies.

### Fitting baseline model to experimental data

For running a single enhancer-gene pair analysis on the experimental data, we obtained the 664 previously published enhancer-gene pairs from the Gasperini et al.^13^ paper using information provided in their Supplemental Table 2. Using these 664 previously published enhancer-gene pairs, we retrieved all experimental gRNAs targeting these enhancers, and filtered gRNAs where there was no valid guide efficiency from GuideScan 2.0 (Table S12). We then obtained the preparation batch, cell gRNA count, and percent mitochondrial reads covariates from their experimental data published on GEO and excluded cells without the ‘guide_count’ covariate value from our downstream modeling. To account for sequencing depth, we used the at-scale gene expression matrix and counted the number of transcripts per cell. We then divided these values by 1e-6 to obtain values for each cell which we included in our linear model through the offset() function. Prior to running the models, a pseudocount of 0.01 was added to the scRNA-seq counts for each cell. Models were then fitted using the nb.glm() function in the MASS R package using a log-link function and optimizing via maximum likelihood estimation. In the at-scale model, there were 207,324 cells total. After filtering for cells without guide count values, there were 205,797 cells that were included in the modeling process. The scrambled perturbation negative control was obtained by scrambling the vector of guide efficiencies prior to modeling. The mismatch gene negative control set was obtained by randomly sampling a gene for a given enhancer from the set of 664 previously published enhancer-gene pairs. Models were successfully run for 609 of the 664 enhancer-gene pairs.

### Defining a model for an enhancer pair acting on a single target gene

Our model for an enhancer gene is quite similar to our baseline model, except we replace **β*_enhancer_* with three new coefficients: *β*_*A*_, *β*_*B*_, *β*_*AB*_. Referring to the two enhancers in the pair being modeled as enhancers A and B: *β*_*A*_represents the effect of enhancer A on the target gene; *β*_*B*_ represents the effect of enhancer B on the target gene; *β*_*AB*_ represents the interaction effect between enhancers A and B on the target gene. The new negative binomial GLM has a mean defined as: *µ* = *exp*(*β*_-_ + **β*_A_X_A_* + **β*_B_X_B_* + **β*_AB_X_AB_* + *β*_*S*_*X*_*S*_ + *β*_*G*2*M*_*X*_*G*2*M*_ + *β*_*m*i*to*_*X*_*m*i*to*_ + *β*_g*RNAs*_*X*_g*RNAs*_ + *β*_*batc*h_*X*_*batc*h_+ *ln*(*s*)) . Here, *X*_*A*_, *X*_*B*_, *X*_*AB*_ represent the perturbation probabilities of enhancer A, enhancer B, and both enhancers, respectively. They are calculated in the same manner as *X_perturb_*.

When fitting linear models, we observed inflated *β*_*AB*_ coefficients associated with cases where all cells containing gRNAs for both enhancers A and B showed no expression of the target gene (Fig. S7). To prevent this inflation of the coefficients, we added a pseudocount of 0.01 to all the gene expression counts. When including a pseudocount in our modeling process, we observed a reduction in outliers in our enhancer effect sizes (Fig. S7).

### Defining testable pairs of enhancers

From the at-scale screen, enhancers and genes were defined using the ‘GSE120861_gene_gRNAgroup_pair_table.at_scale.txt’ file from Gasperini et al., which defined the gRNAgroup-gene pairs tested in their study. To filter for enhancer-gene tests only, gRNAgroup-gene pairs belonging to the ‘NTC’, ‘positive_ctrl’, and ‘TSS’ general groups were removed from our downstream analysis. This resulted in 5,766 unique enhancer sequences and 18,389 unique genes across the gRNAgroup-gene pairs.

The positions of both enhancers and genes were computed as the average between the start and end coordinates. Filtering for enhancers within 1 MB of a gene resulted in 131,356 candidate enhancer-gene links. Some genes had the same Ensembl ID but different positions. After removing these duplicates by keeping the first entry in the table, there remained 128,918 enhancer-gene links. Taking pairwise combinations of these enhancers resulted in 795,616 total enhancer-enhancer-gene sets.

The 795,616 total enhancer-enhancer-gene sets spanned 16,189 unique genes. However, only 9,601 of these genes had gene expression measurements in the scRNA-seq matrix from the at-scale screen. Filtering for enhancer-enhancer-gene sets with complete information yields 477,994 enhancer-enhancer-gene sets.

Enhancer-enhancer-gene sets were then filtered to remove any where both enhancers were not perturbed in at least 10 cells in the study. This was determined as a non-zero count in the matrix of gRNA assignments, which can be found in the ‘GSE120861_at_scale_screen.phenoData.txt.gz’ file from the Gasperini et al. study. Filtering based on this criteria resulted in 82,314 enhancer-enhancer-gene sets.

When computing guide efficiencies to run GLiMMIRS, there is a fraction of guides that do not have efficiency values in the GuideScan database, mostly due to having multiple genome matches or multiple off-target effects (see: Computing guide efficiencies). We removed these guides from downstream analysis, resulting in some enhancers where none of their targeting guides have valid guide efficiency information. Due to this dropout effect, there emerged additional cases where the number of cells with a non-zero perturbation probability for both enhancers is less than 10 which we also filtered out. Ultimately, we were left with 46,166 testable enhancer pairs and corresponding target genes that we successfully ran GLiMMIRS-int on.

### Defining testable pairs of enhancers for high-confidence set

The previously published 664 enhancer-gene pairs were derived from Supplementary Table 2 in the Gasperini paper. Taking pairwise combinations of enhancers that target the same gene from these 664 enhancer-gene pairs resulted in 330 enhancer-enhancer-gene sets. No distance metric limitation was imposed for the high-confidence enhancer-enhancer-gene sets. However, a handful of guide RNAs did not have efficiencies, and were discarded. As a result, we were only able to run GLiMMIRS-int on 264 out of these 330 sets. Not all enhancer pairs in the high-confidence set were jointly perturbed in a minimum of 10 cells (the threshold for defining the larger set of testable enhancer pairs).

### Simulating data for enhancer pairs acting on a single target gene

We adapted the simulation framework used for simulating data for a single enhancer acting on a single gene to simulate data for pairs of enhancers acting on a single gene. We added additional parameters to determine the number of “reference” enhancer pairs with and without an interaction effect between them. We refer to these as “interacting” (*N*_i*nt*_) and “non-interacting” (*N*_*non*_) pairs, respectively. These are selected from the set of all possible pairwise combinations of *N* target sites defined for our simulation. Note that for the case of an enhancer pair acting on a single gene, *N* represents the total number of putative enhancers rather than the total number of target genes. After randomly selecting *N*_i*nt*_ and *N*_*non*_ pairs without replacement from the set of possible pairs, we then randomly select the same number of genes without replacement from the set of possible genes (1, . . . , *G*) to be the target genes of those pairs. For the simulation described in this paper, we selected values of *N*_i*nt*_ = *N*_*non*_ = 500 and a total of *N* = 1000 target sites.

### Simulating data for power analysis

Most aspects of the data simulation are identical to the data simulation for a single enhancer acting on a single gene. The coefficients *β*_*A*_and *β*_*B*_are drawn from a normal distribution defined as *N*(*µ* = −0.02, *σ* = 0.16). The cell cycle effects *β*_*S*_ and *β*_*G*2*M*_ were also sampled from normal distributions defined as *N*(*µ* = −0.21, *σ* = 1.04) and *N*(*µ* = 0.004, *σ* = 0.49), respectively. The effect size for percentage of mitochondrial DNA, *β*_*m*i*to*_, was sampled from a normal distribution defined as *N*(*µ* = −0.37, *σ* = 9.04). These distributions were selected by fitting to the enhancer effects estimated from the experimental data. For the power analysis, we assign a number of different fixed values of *β*_*AB*_for genes that are acted upon by an interaction effect between enhancers (e.g., the target genes of “positive” enhancer pairs). For genes that are not acted upon by any interaction effect, we set *β*_*AB*_ = 0. The other parameter that we modulate in the simulations is the value of λ for the Poisson distribution used to sample the number of unique gRNAs found in each cell. This is representative of multiplicity of infection, or MOI, so for each value of λ that we want to test with our power analysis, we generate different numbers of gRNAs per cell and use these sets of values to generate different mappings of gRNAs in cells. This yields a different one-hot encoded matrix for each value of lambda, which will also lead to different sets of values of *X*_*A*_, *X*_*B*_, and *X*_*AB*_, as greater MOI may result in more gRNAs for a target site found in each cell and greater perturbation probabilities. Simulated counts are generated from a negative binomial distribution parameterized by *NB*(*µ*, *ϕ*), where *µ* = *exp*(*β*_-_ + **β*_A_X_A_* + **β*_B_X_B_* + **β*_AB_X_AB_* + *β*_*S*_*X*_*S*_ + *β*_*G*2*M*_*X*_*G*2*M*_ + *β*_*m*i*to*_*X*_*m*i*to*_ + *ln*(*s*)) and *ϕ* = 1.5 (determined from modeling empirical data, see Methods for simulating data for single enhancers acting on a single gene). We generated a set of simulated counts for each value of λ and interaction effect size. For our power analysis, we used values of λ = 15,20, 30, 50 and *β*_*AB*_ = 0.5,1,2,3,4,5,6.

### Power analysis

For our power analysis, we fit our model to the simulated data for the “positive” and “negative” pairs to obtain true positive rates (TPR) and true negative rates (TNR), respectively. We calculated the proportion of models that correctly called significant interaction terms, *β*_*AB*_, for the “positive” cases to obtain TPR. We calculated the proportion of models that correctly called no significant interaction terms, *β*_*AB*_, for the “negative” cases to obtain TNR.

### Comparing multiplicative to additive model

To compare the fits of multiplicative vs. additive models of enhancer pair activity, we defined each model under the null hypothesis (*H*_0_), where there is no interaction term (for simplicity). For the multiplicative model under *H*_0_, we use the canonical log-link function and define the mean of the negative binomial, *µ*, as: ^*µ* = *exp*(*β*^_0_ ^+ *β*^_*A*_^*X*^_*A*_ ^+ *β*^_*B*_^*X*^_*B*_ ^+ *β*^_*S*_^*X*^_*S*_ ^+ *β*^_*G*2*M*_^*X*^_*G*2*M*_ ^+ *β*^_*m*i*to*_^*X*^_*m*i*to*_ ^+ *β*^_g*RNAs*_^*X*^_g*RNAs*_ ^+ *β*^_*batc*h_^*X*^_*batc*h_ ^+^ *ln*(*s*)). For the additive model under *H*_0_, we use the identity link function where the mean is simply equivalent to the linear predictor without transformation, defined as: ^*µ* = *s*(*β*^_-_ ^+ *β*^_*A*_^*X*^_*A*_ ^+ *β*^_*B*_^*X*^_*B*_ ^+ *β*^_*AB*_^*X*^_*AB*_ ^+ *β*^_*S*_^*X*^_*S*_ ^+ *β*^_*G*2*M*_^*X*^_*G*2*M*_ ^+ *β*^_*m*i*to*_^*X*^_*m*i*to*_ ^+ *β*^_g*RNAs*_^*X*^_g*RNAs*_ ^+^ *β*_*batc*h_*X*_*batc*h_). We applied each model to the 330 testable pairs from the experimental data where each enhancer in the pair had evidence of being an enhancer for the target gene based on the analysis by Gasperini et al. We compare model fits by examining the Akaike Information Criterion (AIC), with a lower AIC indicating a better fit. We calculated *ΔAIC* by subtracting the AIC of the lesser model from the AIC of the best fitting model. Since we found that the multiplicative model fit better in every case we tested, every *ΔAIC* reported in our study reflects the AIC of the additive model subtracted from the AIC of the multiplicative model.

### Fitting interaction model to empirical data

For analyzing both sets of enhancer pairs tested in our analysis, we followed an identical procedure to the baseline model scenario, with the exception of adding a second enhancer effect vector, and allowing for interactions between the two enhancer vectors using built-in functionality within the glm.nb() function in the MASS R package.

### Cook’s Distance Outlier Filtering

When analyzing the significant enhancer-enhancer interactions from our at-scale analysis, we observed that several of these interactions were large in magnitude and positive. We then manually inspected the gene expression counts of cells with both enhancers perturbed and observed that this cell population generally had a single cell with a high expression. Since coefficient estimates from generalized linear models can be strongly influenced by outliers, we believe the “significant” interactions detected by GLiMMIRS-int in the Gasperini analysis are artifacts of the single-cell CRISPR experiment rather than true biological signals. To identify outlier cells that drive interaction coefficient estimates, we used Cook’s distance, which is a metric that quantifies the influence of each observation on the coefficients of a regression model and which has been previously used in differential expression analysis tools like DESeq2^44^. To define outlier-driven interactions, we set a threshold of: 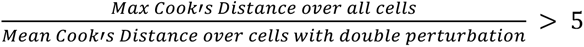, and we discarded enhancer-enhancer interactions with jointly perturbed cells that met this criterion from downstream analysis. We applied this same metric to the significant interactions detected in the Morris et al. dataset.

### Bootstrap expression confidence intervals

We used a bootstrapping procedure to compute confidence intervals for gene expression estimates and the expected expression under a multiplicative model (Fig. 3E,F). For each of the enhancer-enhancer interactions that remained significant in our at-scale analysis following our Cook’s distance-based filtering, we performed 100 bootstrap sampling iterations, in which all cells were resampled with replacement. In each iteration, the full GLiMMIRS-int model was fit to the data, and the intercept, enhancer 1 perturbation (E1), enhancer 2 perturbation (E2), and interaction coefficient estimates were recorded. Using these coefficients, scaled expression values were computed using the following formulas:

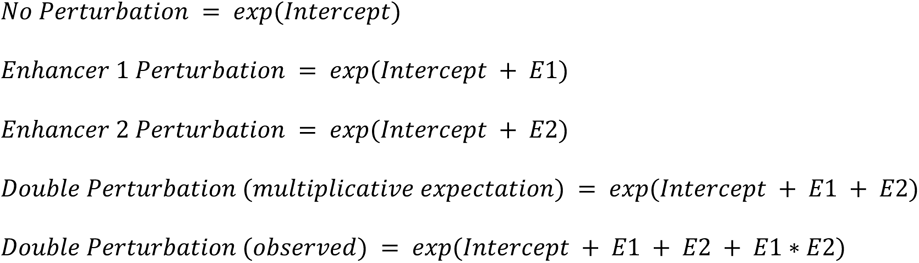

The central 90 coefficient estimates from the bootstrap iterations were used to compute 90% confidence intervals.

### Preprocessing Morris et al. CRISPRi data for *PTPRC* Analysis

Single-cell RNA-sequencing and guide RNA data were downloaded from GEO accession number GSE171452 under ‘STINGseq-v1_cDNA’ (<u>GSM5225857</u>) and ‘STINGseq-v1_GDO’ (<u>GSM5225859</u>), respectively. Cell-level covariates were obtained from Supplementary Table S3C. SNP IDs for the SNPs surrounding the *PTPRC* locus were obtained from Supplementary Table S1E. SNP-guide RNA assignments were obtained from Supplementary Table S3A. All supplementary tables referenced were downloaded directly from the Morris et al. manuscript^28^.

Cell cycle scoring was performed on the scRNA-sequencing data using the Seurat software package. Cells were pre-filtered using the same QC metrics as Morris et al. Cell cycle scores were computed using the following functions: Read10X(), CreateSeuratObject(),

NormalizeData(), FindVariableFeatures() with the ‘vst’ selection method, ScaleData() with all genes in the RNA-sequencing matrix, and CellCycleScoring() using the predefined S and G2M gene sets from Seurat. All functions were run with default parameters unless otherwise specified.

The gRNA assignment matrix was binarized based on whether each count value was greater than or equal to its gRNA UMI threshold. Guide RNA UMI thresholds were determined using the ‘umi_thresholds.csv.gz’ file from GEO. For the STING-seq analysis, we did not include guide efficiency information, since none of the guide RNA sequences targeting *PTPRC* enhancers in the experiment had efficiencies available in the GuideScan database.

Prior to running GLiMMIRS, both the guide RNA and RNA-sequencing matrices were pre-filtered to only include cells passing the QC thresholds previously defined by Morris et al.

### In silico perturbation experiments with Enformer

To select enhancer pairs for input into Enformer, we first filtered the set of 795,616 enhancer pairs in the Gasperini data set to those that were simultaneously perturbed in a minimum of 10 cells. We then reduced this set to enhancer pairs associated with expressed genes in our single-cell RNA-sequencing matrix. To evaluate the effects of synthetic perturbations on a target gene of interest, we focused on enhancer pairs that were both located in a ∼196kb window centered on the TSS of their putative target gene as these would be possible to evaluate with Enformer given the model’s input size constraints (196,608bp). These criteria reduced the set of enhancer pairs to 2,136 pairs. We then retrieved the input sequences containing each of these enhancer pairs from hg19 and used Enformer to make predictions on these wild type (WT) unperturbed sequences. Enformer predicts CAGE-seq reads over a region of 114,688bp at the center of the input sequence in 128 bp bins, resulting in 896 output bins altogether. To focus on TSS activity only, we considered the average of the predictions for the central bins (i.e. bins 447 and 448) across 10 different shuffles of the enhancer regions as the predicted gene expression signal for a given gene. We used predictions for track 5,111 of the human output head, which corresponds to K562 CAGE predictions. We then removed sequences where the Enformer predicted WT signal was below 10, leaving a total of 1372 sequences for downstream analysis. We then made predictions with the first enhancer in the pair shuffled (EnhA); the second enhancer in the pair shuffled (EnhB); and both enhancers in the pair shuffled (EnhA&EnhB). Shuffles were accomplished by performing ten independent dinucleotide shuffles of the target enhancer(s) and averaging Enformer’s predictions, similar to a Global Importance Analysis^45^. This shuffling and averaging serves to marginalize out the contribution of the enhancer(s), similar to inactivation perturbation via CRISPRi. The difference between the average predicted CAGE-seq level between mutant (i.e. shuffled) sequences and WT sequences was quantified with a log ratio: 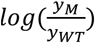, where y_m_ is the Enformer-predicted expression of the mutant sequence and *y*_*WT*_ is the Enformer-predicted expression of the wild type sequence. We estimated enhancer effects for the individual mutant sequences under a multiplicative model as:

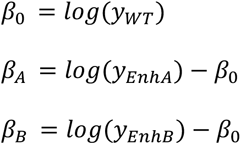

and then computed the expected expression of the target gene for the double mutant sequence under the multiplicative model as:

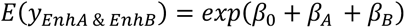

We then plotted the expected difference in expression 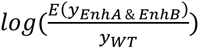 against the Enformer-predicted (observed) difference in expression 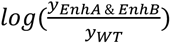 and calculated the Pearson correlation between them (Fig. 4C).

### Bayesian Interaction Detection Probability Analysis

We performed a Bayesian analysis to obtain posterior estimates of the frequency of enhancer interactions, given different prior beliefs in interaction frequencies. We used a beta distribution to specify priors for interaction frequency, with means between 0.05 and 1, and standard deviation of 0.2. Then, we considered various degrees of power to detect interactions: 0.05, 0.1, 0.2, 0.4, 0.6, 0.8. For each power setting, we computed a prior probability distribution for detecting enhancer-enhancer interactions using GLiMMIRS by scaling the mean and standard deviation estimates by the power. Since the prior is a beta distribution and the number of detected interactions follows a binomial distribution, the prior is conjugate and the posterior distribution is also a beta distribution with a closed-form solution. The curves in Fig. 3G and 3H show the maximum a posteriori probability (MAP) from the resulting posterior distribution. For the high-confidence set of enhancer pairs (Fig. 3G), we set the number of detected interactions to 0 out of 264 tested. For the entire set of enhancer pairs (Fig. 3H), we excluded the significant outlier-driven interactions and set the number of detected interactions to 11 out of 46,166 tested.

## Data and code availability

● Data from the Gasperini et al. experiment can be found under GEO accession number GSE120861.
● Our *NMU* RT-qPCR experiment results (Table S1) are provided as a spreadsheet.
● All relevant code and documentation can be found at https://github.com/mcvickerlab/GLiMMIRS.

## Acknowledgements

We thank the research groups of Dr. Roland Schwarz and Dr. Christoph Lippert for hosting J.L.Z. during her time in Berlin as a visiting Fulbright scholar and supporting her work on this project. We thank John Morris for assistance with the Morris et al. dataset. We thank the anonymous reviewers for their constructive criticism of the manuscript. We thank Wilfred Wong for assistance with visualizing Hi-C data in the WashU epigenome browser. J.L.Z. was supported by a Fulbright Research Award, an NIH F31 individual predoctoral fellowship (F31DA056226), and the Chapman Charitable Trust Fellowship. G.M. was supported by the NIH/NHGRI (R35HG011315) and the Frederick B. Rentschler Developmental Chair. H.V.C. was supported by the 2020 Salk Women & Science Award and the 2020 Salk Alumni Fellowship Award.

## Author Information

### Author contributions

J.L.Z. and K.G. contributed equally as co-first authors and have agreed that either author can be listed first in reporting this study. The project was conceptualized by G.M. and J.L.Z. J.L.Z. and K.G. performed computational analyses, data curation, and methods development. J.L.Z. developed the data simulation pipeline and performed all analyses on the simulated data, including the power analysis. K.G. worked on GLiMMIRS-base and GLiMMIRS-int and applied them to the Gasperini et al. and Morris et al. datasets. K.G. also performed outlier detection and the Bayesian analysis for frequency of enhancer interactions. J.L.Z., G.M., and K.G. wrote the manuscript. S.T. performed the in-silico perturbation experiment with Enformer and J.L.Z. analyzed the results. P.K. advised S.T. H.V.C and A.R.C. performed the knockout experiments on the *NMU* locus. P.K., S.T., and H.V.C. provided feedback on the manuscript.

---

**Figure S1.**
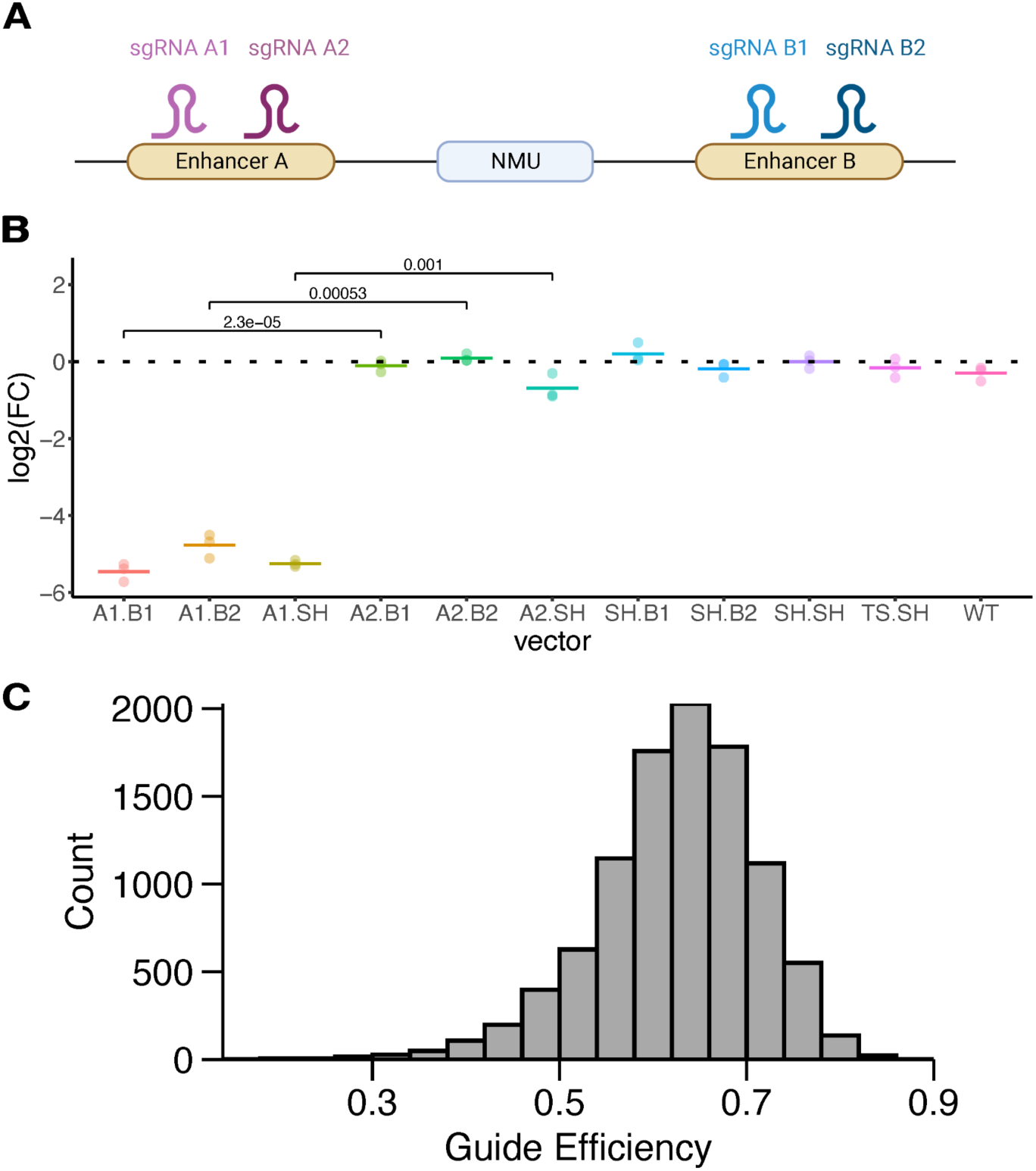
Variable gRNA efficiency may bias interpretation of enhancer interactions. **A)** We examined two enhancers of *NMU*, which were among the most significant enhancer-gene pairs discovered by Gasperini et al. We performed CRISPRi experiments to perturb the enhancers of *NMU* using guide designs from the original study (Table S1). **B)** Results of CRISPRi RT-qPCR experiment perturbing NMU enhancers for three technical replicates. For each NMU enhancer (enhancers A and B), two gRNAs were used (A1, A2 and B1, B2, respectively) and delivered on the same vector. Vectors containing gRNA A1 resulted in larger fold changes in NMU expression than their counterparts containing gRNA A2 instead (denoted p-values come from unpaired Welch’s two-sided t-tests against the null hypothesis that there is no difference in mean fold change (FC) between vectors using gRNA A1 vs. gRNA A2. SH = safe harbor). TS = NMU transcription start site, WT = wild type K562 cells expressing dCas9-KRAB without any gRNAs, horizontal bar = mean log_2_(FC). See also Table S1. **C)** Distribution of guide efficiency values predicted by GuideScan 2.0 for the gRNAs used in the Gasperini et al. experiment.

**Figure S2.**
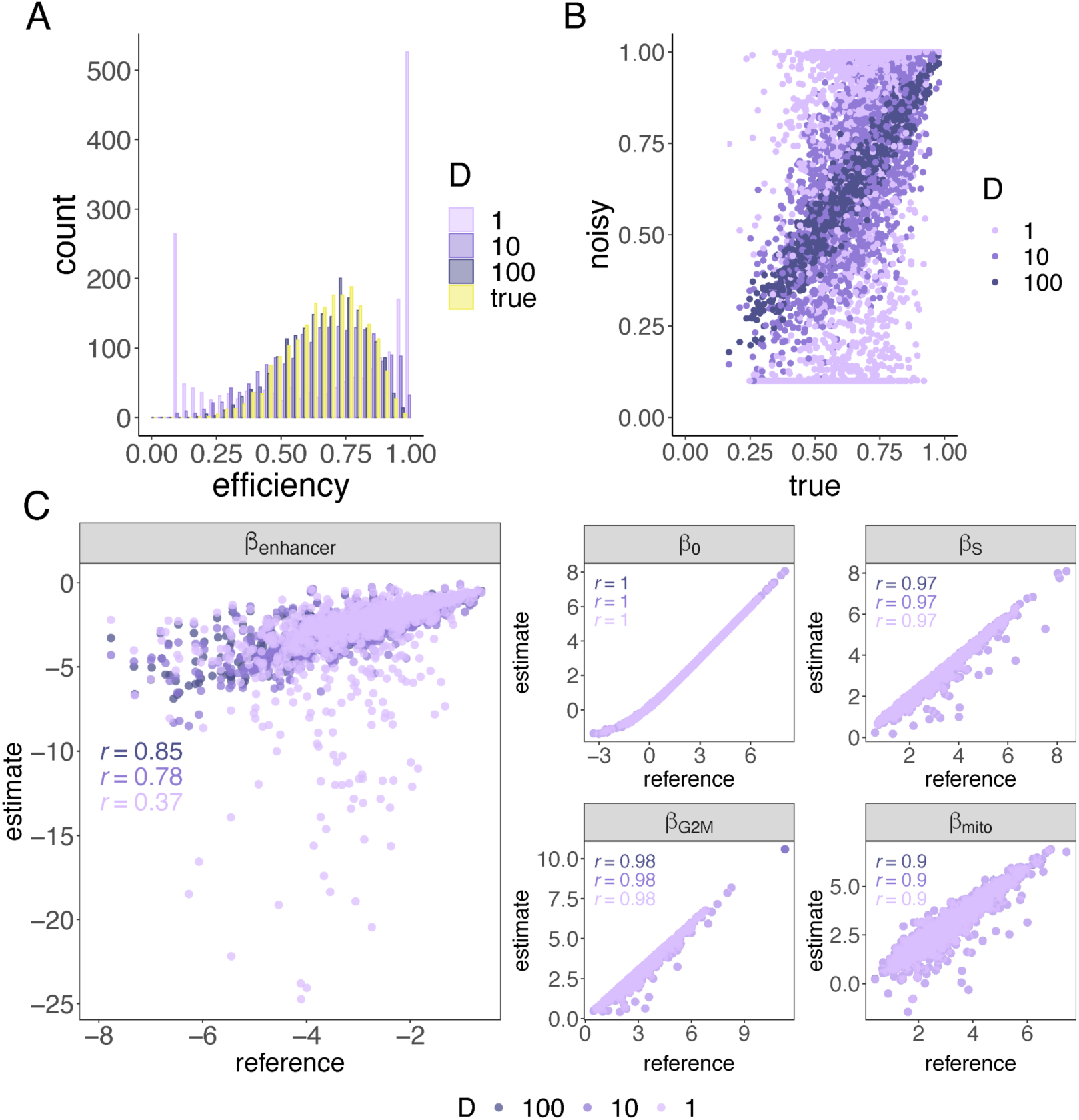
Simulated noisy gRNA efficiency values and their effects on coefficient estimates. **A)** Histogram of noisy and true guide efficiencies from simulations with different values of *D*, the dispersion-controlling coefficient used to control “noise.” **B)** Scatterplot comparing noisy guide efficiencies to true guide efficiencies with different values of *D*. Pearson r = 0.391, 0.716, 0.956 for D=1, 10, 100, respectively. **C)** Scatterplot comparing true versus estimated coefficient values for each gene evaluated with GLiMMIRS-base. These plots summarize the results of fitting the model to 1000 genes in the simulated dataset which were designated as “true” target genes (genes whose enhancers were perturbed by gRNAs in the simulated experiment). Plot shows results of fitting to simulated data using the three different sets of noisy guide efficiencies. A pseudocount of 0.01 was applied to the counts for all cells. Pearson correlations (*r*) are shown here and in Table S3. 36 outliers fall outside the axis range and are not visible in the **β*_enhancer_* panel for the set where D=1.

**Figure S3.**
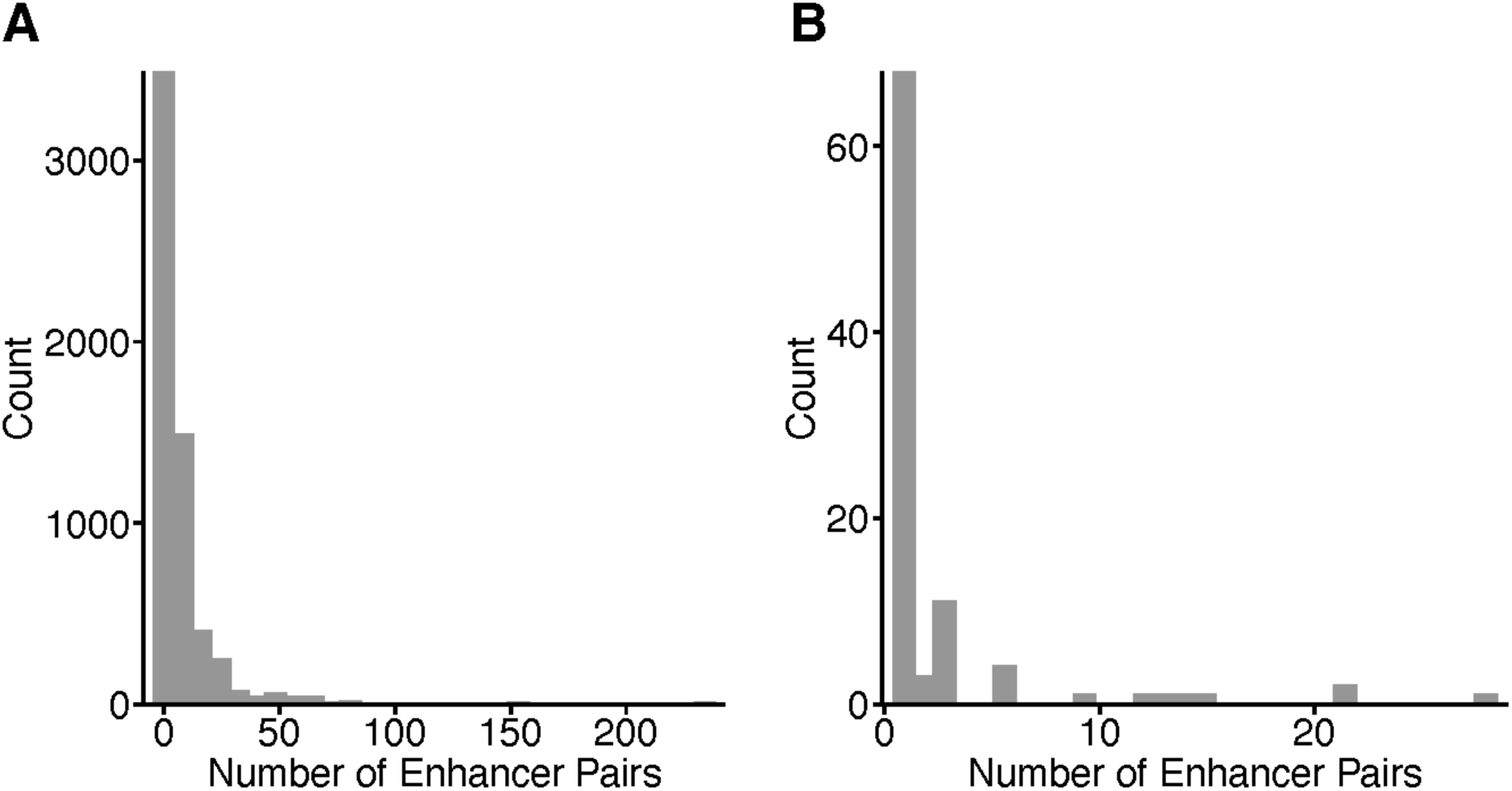
Number of enhancer pairs per gene. For **A)** the entire testable set of 46,166 enhancer pairs; and **B)** the “high confidence” set of 264 enhancer pairs.

**Figure S4.**
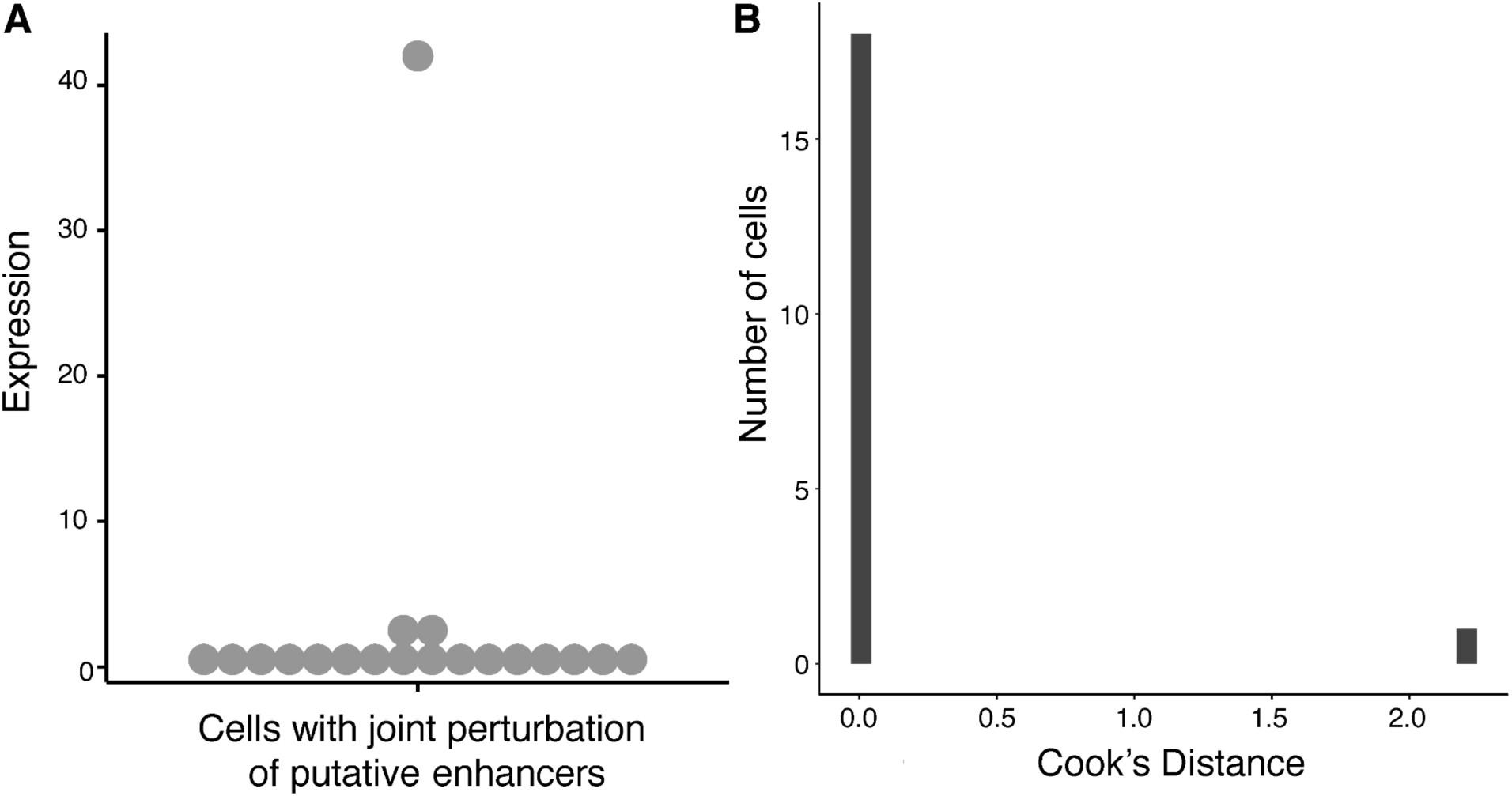
Example of enhancer pair with outlier gene expression. **A)** Expression of gene *BABAM2* in cells with jointly perturbed enhancers. **B)** Distribution of Cook’s distances for the same cells.

**Figure S5.**
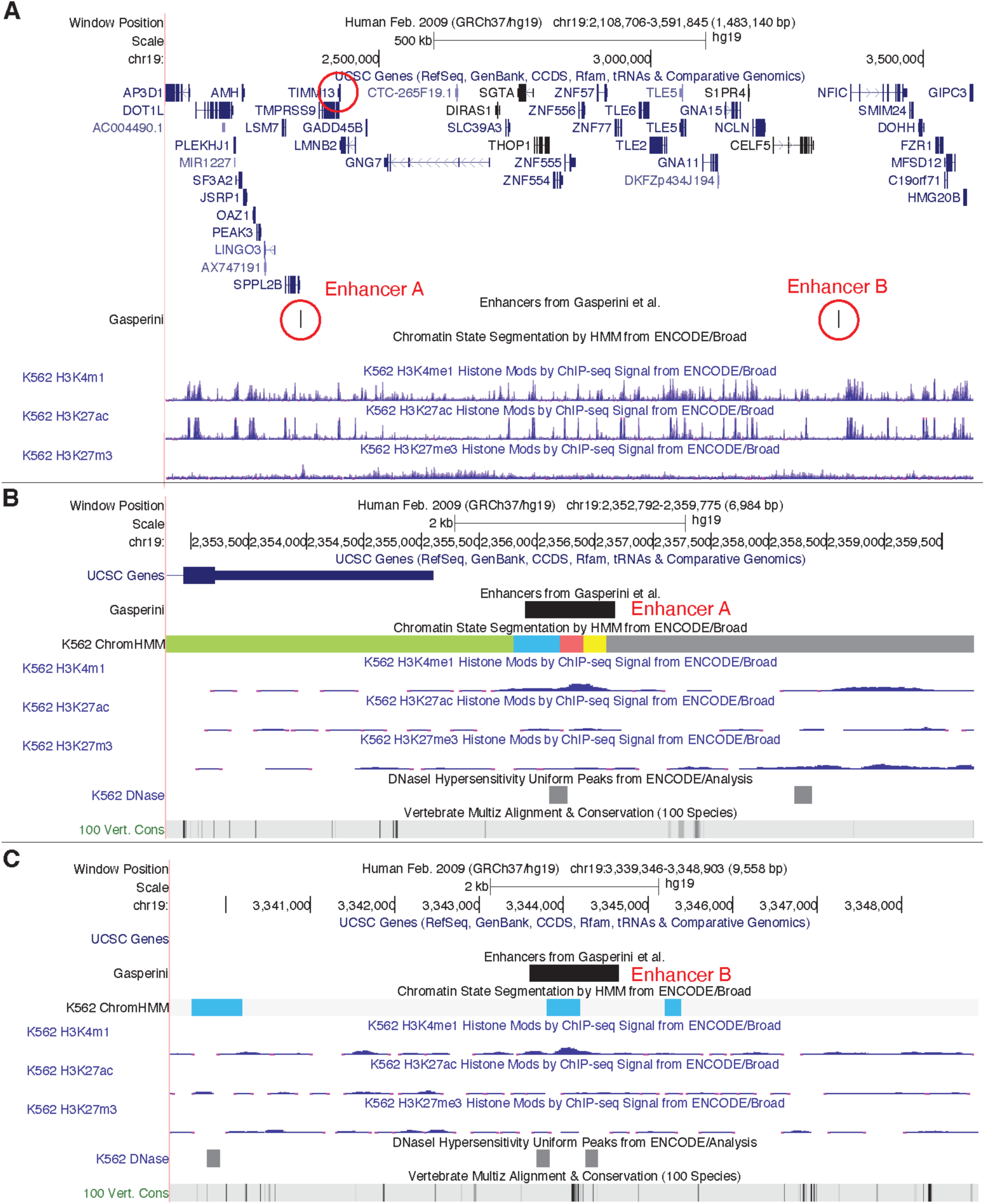
A pair of enhancers with a negative interaction effect on the expression of *TIMM13*. **A)** Overview showing entire genome regions and both enhancers. **B)** Zoom-in of enhancer A region. **C)** Zoom-in of enhancer B region.

**Figure S6.**
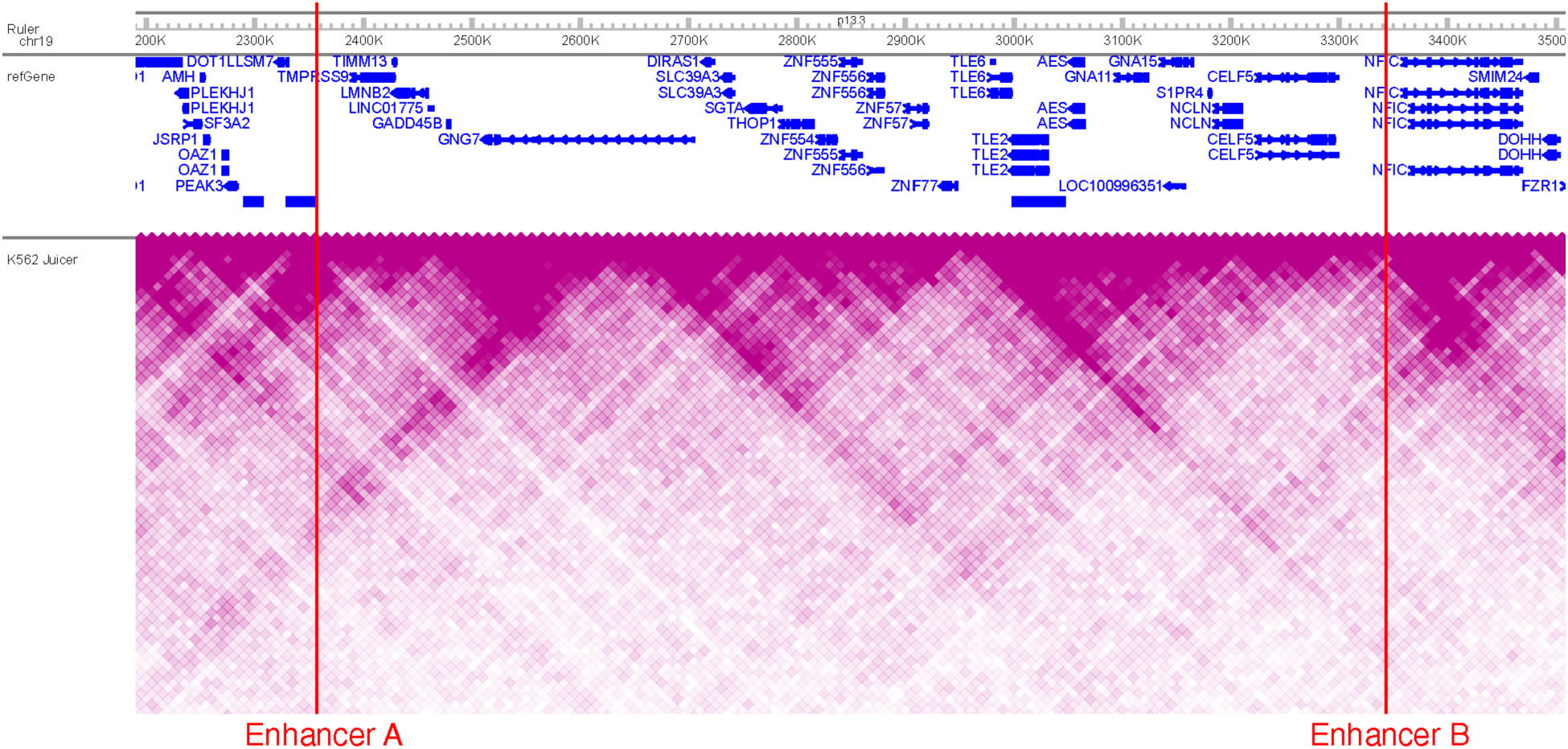
Enhancers with a negative interaction effect on the expression of TIMM3 are in different topological associating domains (TADs). The heatmap shows the Hi-C contact frequency in K562 cells^46^ from the WashU epigenome browser^47^ and the locations of both enhancers.

**Figure S7.**
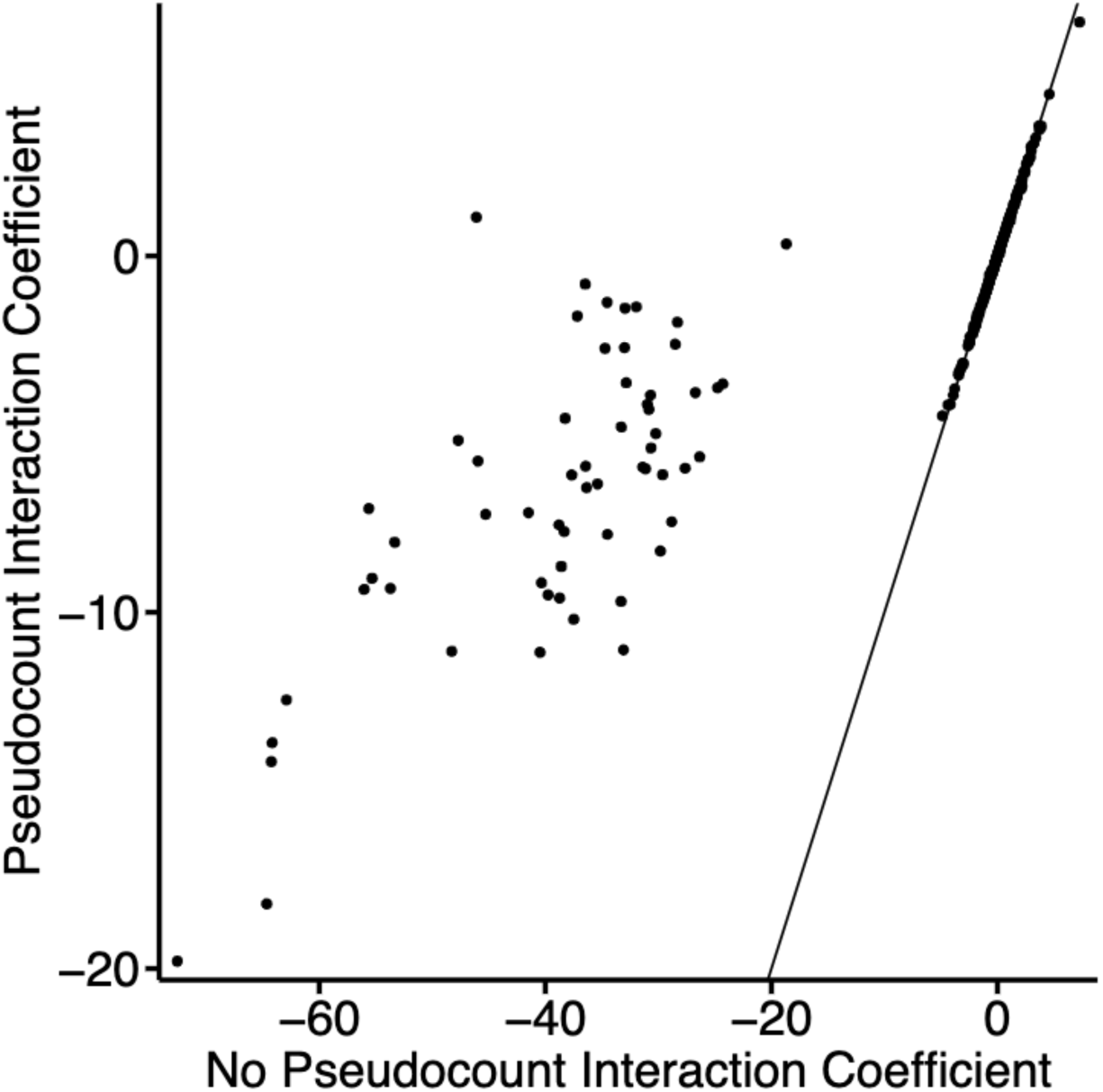
Outlier interaction coefficient estimates are moderated by introduction of a pseudocount. Magnitude of interaction term coefficients for 330 enhancer-enhancer pairs when adding vs. not adding a pseudocount of 0.01 to adjust the gene expression. The inclusion of a pseudocount greatly reduces the magnitude of outlier interaction coefficient estimates (note difference in x and y axis scales).

## Supplementary Tables

**Table S1. Data from *NMU* RT-qPCR experiment.** Provided separately as an Excel file.

**Table S2.**
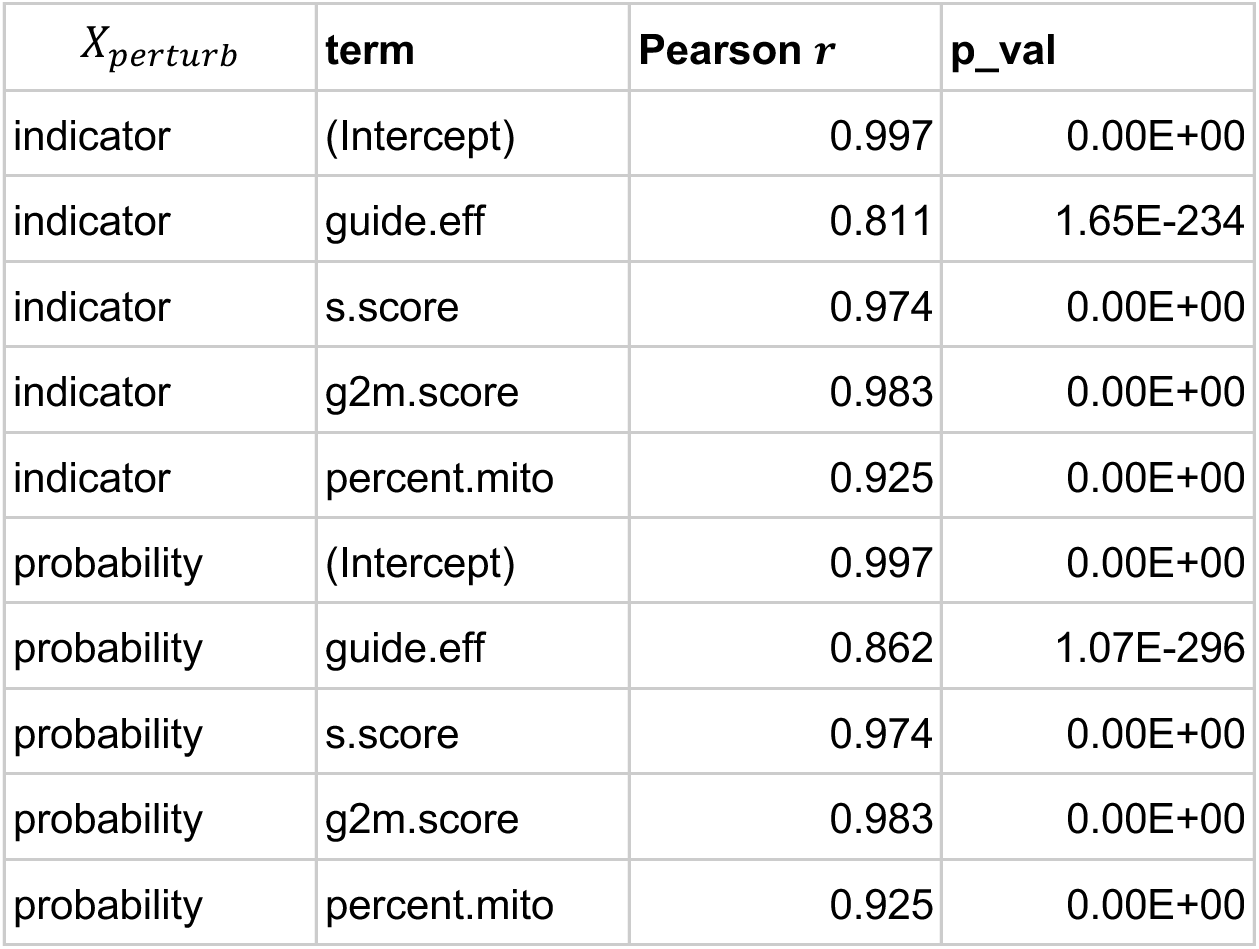
Fitting GLiMMIRS-base to simulated data comparing perturbation probability to indicator variable for X_*perturb*_. Pearson correlation (Pearson *r*, p_val) between reference and estimated coefficient values for each coefficient in the baseline model when fitting with guide-efficiency derived value of *X_perturb_* versus with an indicator (0/1) value for *X_perturb_*.

**Table S3.**
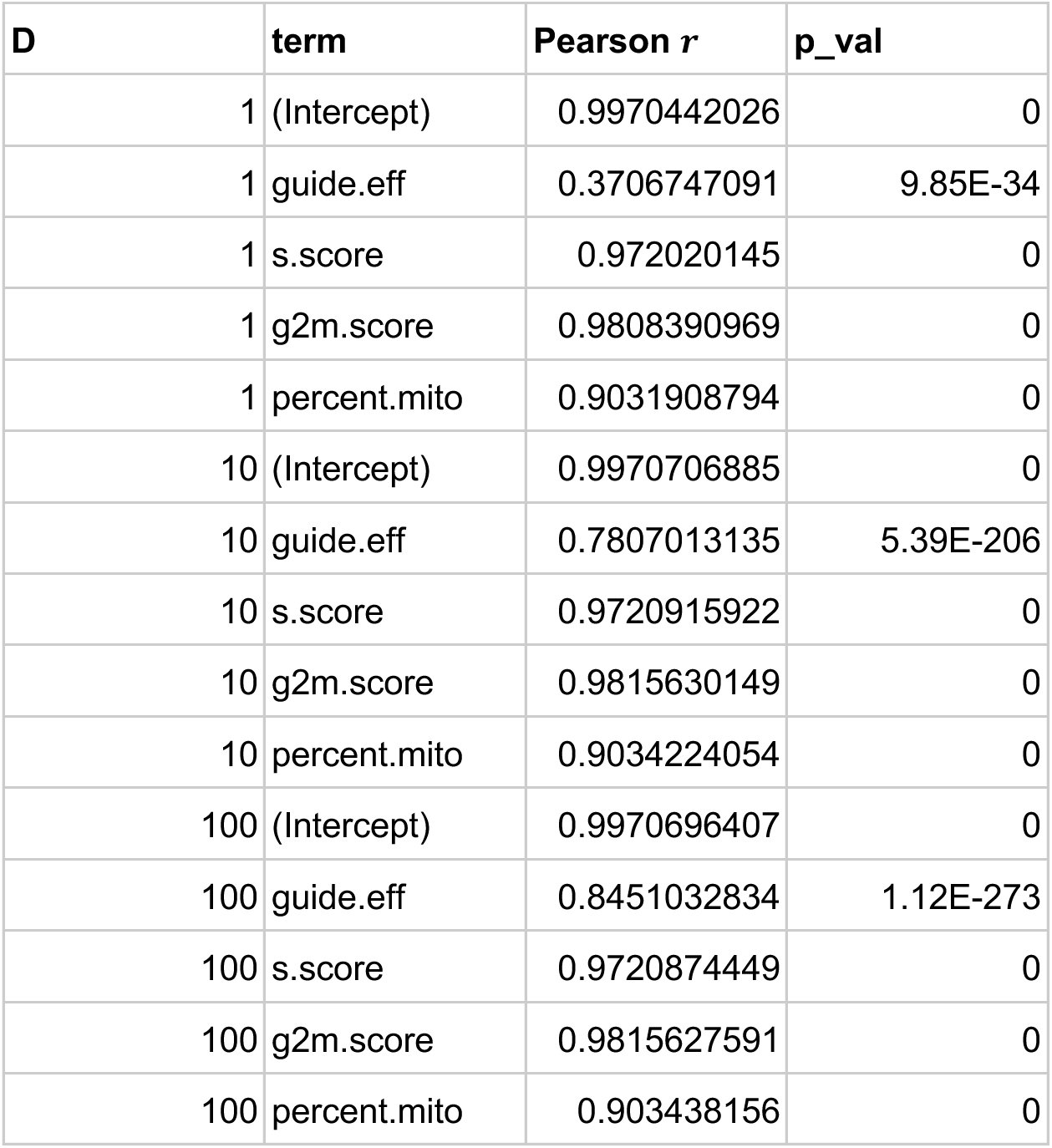
Fitting GLiMMIRS-base to simulated data comparing different levels of noise in guide efficiency estimates. Pearson correlation (*r*), between reference and estimated coefficient values for each coefficient in the baseline model when fitting with perturbation probabilities *X_perturb_* calculated from different sets of noisy guide efficiency estimates (*D* = 1 high noise; *D* = 10 moderate noise, *D* = 100 low noise).

**Table S4. Results from applying GLiMMIRS-base to high-confidence enhancers from Gasperini et al.** (Provided separately as .csv file).

**Table S5. Power analysis for GLiMMIRS-int.** Power analysis results for different values of λ (lambda) and interaction effect sizes (effect.size). True positive rate (TPR) is calculated from simulated data for interacting enhancer pairs and false positive rate (FPR) is calculated from simulated data for non-interacting enhancer pairs. (Provided separately as .csv file).

**Table S6. Results from applying GLiMMIRS-int to high-confidence enhancers from Gasperini et al.** (Provided separately as .csv file).

**Table S7. Results from applying GLiMMIRS-int to complete set of testable enhancers from Gasperini et al.** (Provided separately as .csv file).

**Table S8. Summary of Cook’s distance estimates.** Cook’s distance for the interaction coefficient was computed for each cell with joint perturbations for the 31 enhancer-pairs with significant interaction effects. (Provided separately as .csv file).

**Table S9. Results from applying GLiMMIRS-base to candidate enhancers for *PTPRC* using data from Morris et al. 2023** (Provided separately as .csv file).

**Table S10. Results from applying GLiMMIRS-int to candidate enhancer pairs and *PTPRC* using data from Morris et al. 2023.** (Provided separately as .csv file).

Table S11. Enformer gene expression predictions in the presence of perturbations to individual enhancers and joint perturbations for 2136 enhancer pairs and corresponding target genes tested. (Provided separately as .csv file).

**Table S12. Results from applying GuideScan 2.0 to Gasperini et al. guide sequences.**

(Provided separately as .csv file).

Table S13. Ensembl to HGNC gene mapping used for cell cycle analysis. (Provided separately as .csv file).

Table S14. Cell cycle S scores computed with Seurat. (Provided separately as .csv file).

Table S15. Cell cycle G2M scores computed with Seurat. (Provided separately as .csv file).

