## Supplemental Information for "Analysis of single-cell CRISPR perturbations indicates that enhancers act multiplicatively and provides limited evidence for epistatic-like interactions"

### Supplementary Figures

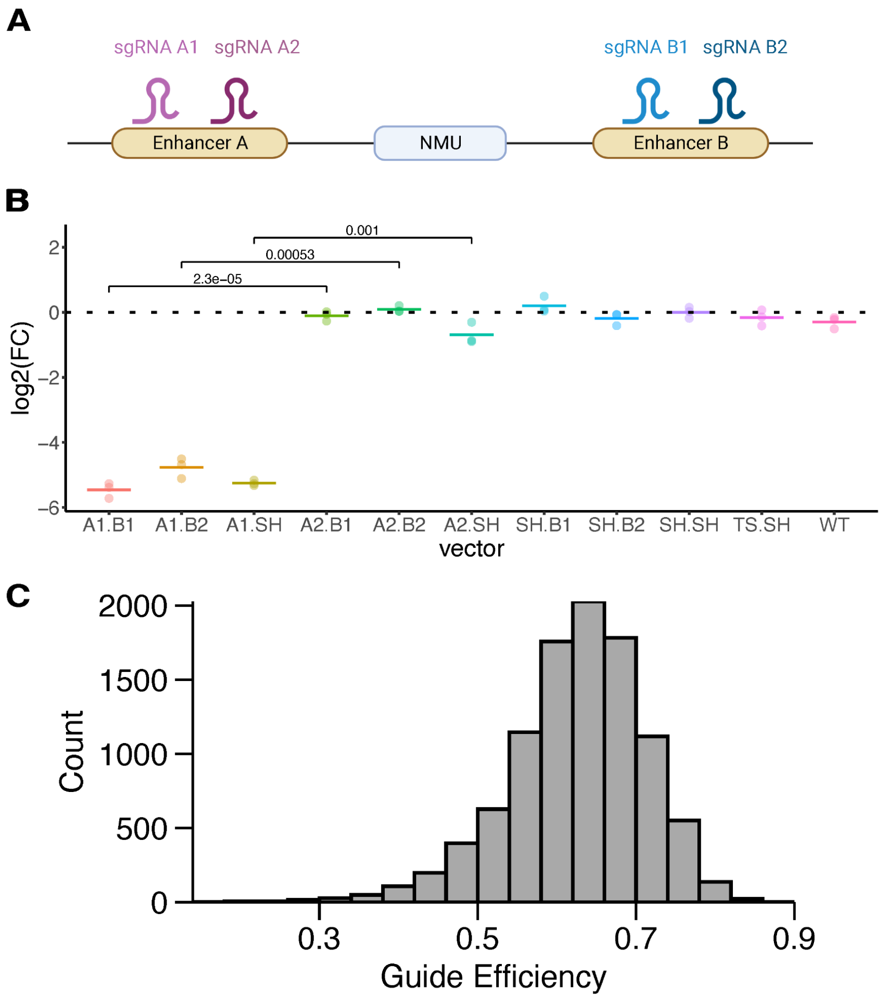


**Figure S1. Variable gRNA efficiency may bias interpretation of enhancer interactions. A)** We examined two enhancers of *NMU*, which were among the most significant enhancer-gene pairs discovered by Gasperini et al. We performed CRISPRi experiments to perturb the enhancers of *NMU* using guide designs from the original study (Table S1). **B)** Results of CRISPRi RT-qPCR experiment perturbing NMU enhancers for three technical replicates. For each NMU enhancer (enhancers A and B), two gRNAs were used (A1, A2 and B1, B2, respectively) and delivered on the same vector. Vectors containing gRNA A1 resulted in larger fold changes in NMU expression than their counterparts containing gRNA A2 instead (denoted p-values come from unpaired Welch's two-sided t-tests against the null hypothesis that there is no difference in mean fold change (FC) between vectors using gRNA A1 vs. gRNA A2. SH = safe harbor). TS = NMU transcription start site, WT = wild type K562 cells expressing dCas9-KRAB without any gRNAs, horizontal bar = mean $\log_2$(FC). See also Table S1. **C)** Distribution of guide efficiency values predicted by GuideScan 2.0 for the gRNAs used in the Gasperini et al. experiment.

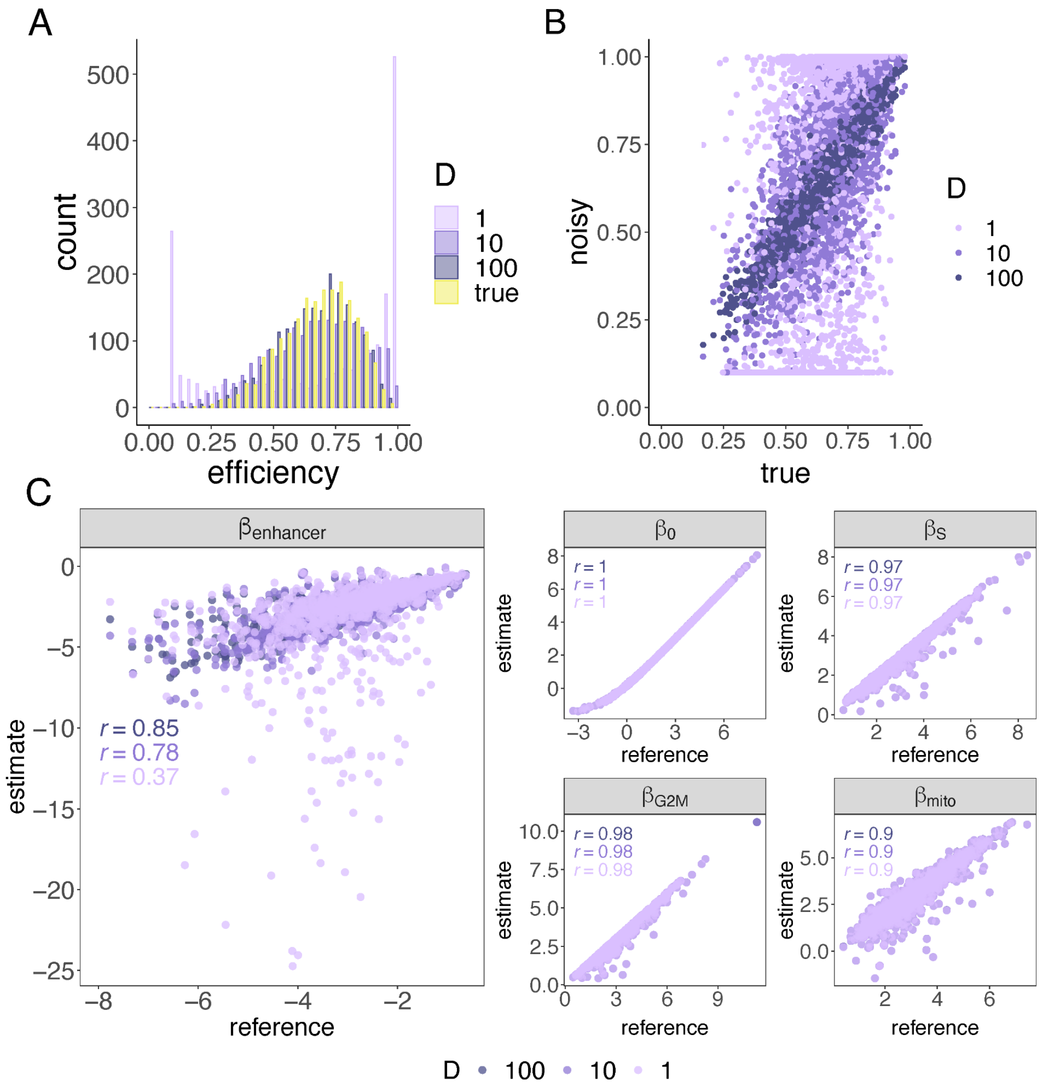


**Figure S2. Simulated noisy gRNA efficiency values and their effects on coefficient estimates. A)** Histogram of noisy and true guide efficiencies from simulations with different values of $D$, the dispersion-controlling coefficient used to control "noise." **B)** Scatterplot comparing noisy guide efficiencies to true guide efficiencies with different values of $D$. Pearson r = 0.391, 0.716, 0.956 for D=1, 10, 100, respectively. **C)** Scatterplot comparing true versus estimated coefficient values for each gene evaluated with GLiMMIRS-base. These plots summarize the results of fitting the model to 1000 genes in the simulated dataset which were designated as "true" target genes (genes whose enhancers were perturbed by gRNAs in the simulated experiment). Plot shows results of fitting to simulated data using the three different sets of noisy guide efficiencies. A pseudocount of 0.01 was applied to the counts for all cells. Pearson correlations ($r$) are shown here and in Table S3. 36 outliers fall outside the axis range and are not visible in the $\beta_{enhancer}$ panel for the set where D=1.

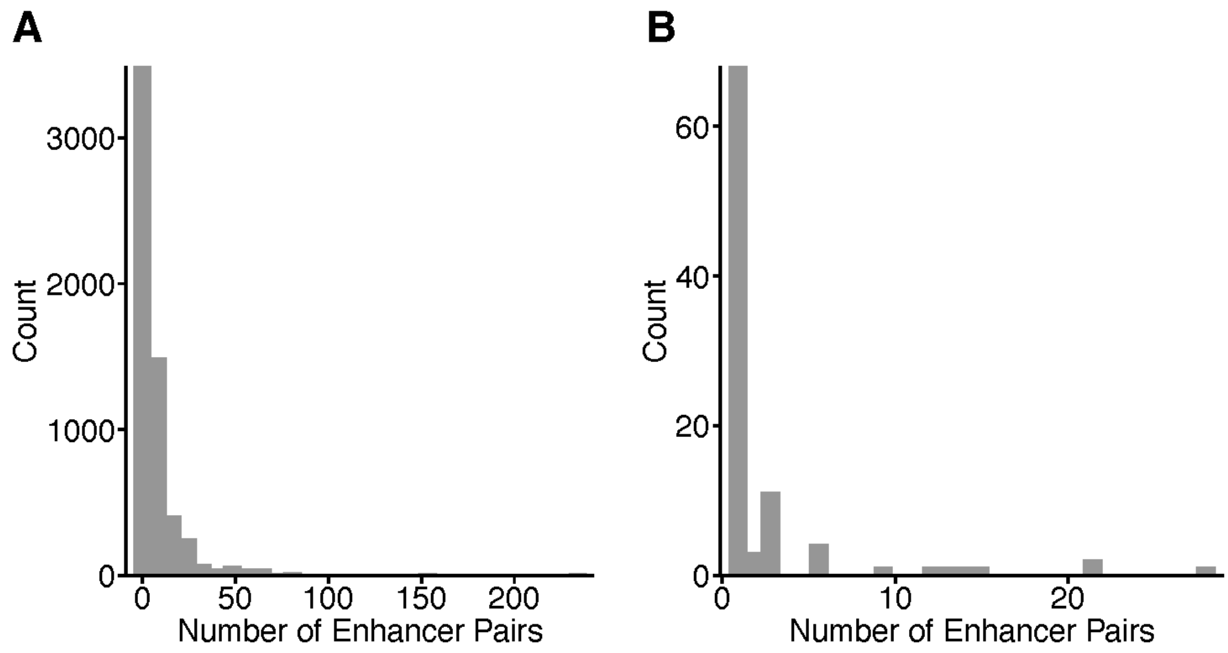


**Figure S3. Number of enhancer pairs per gene.** For **A)** the entire testable set of 46,166 enhancer pairs; and **B)** the "high confidence" set of 264 enhancer pairs.

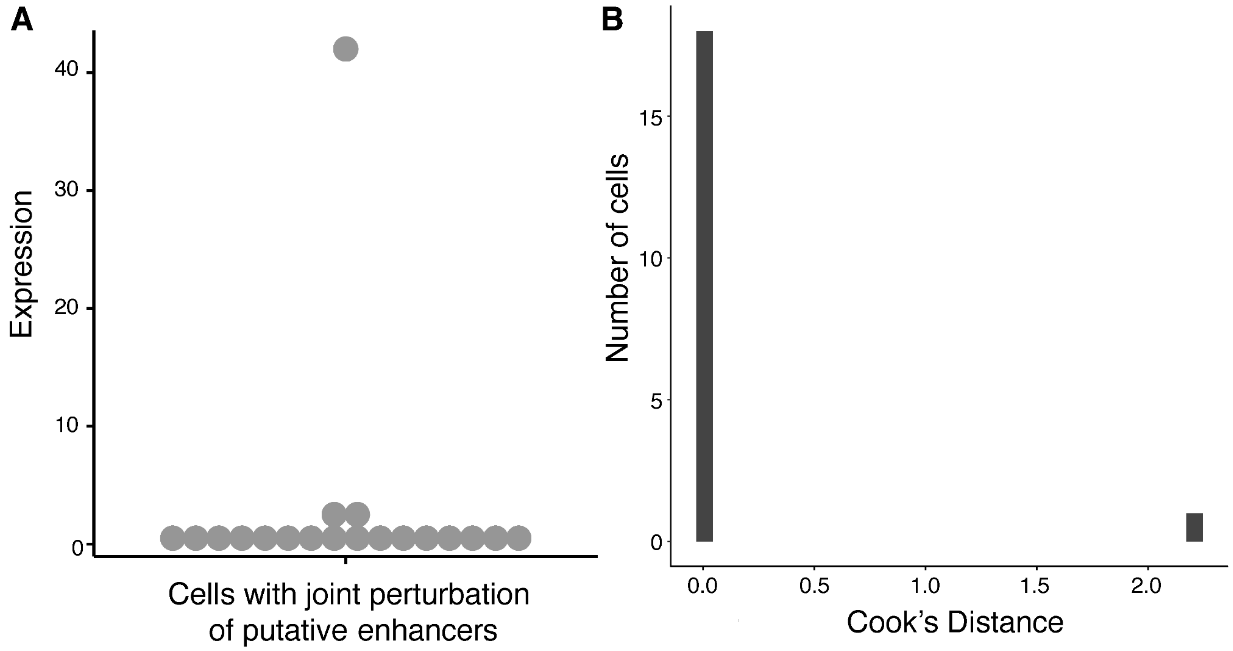


**Figure S4. Example of enhancer pair with outlier gene expression. A)** Expression of gene *BABAM2* in cells with jointly perturbed enhancers. **B)** Distribution of Cook's distances for the same cells.

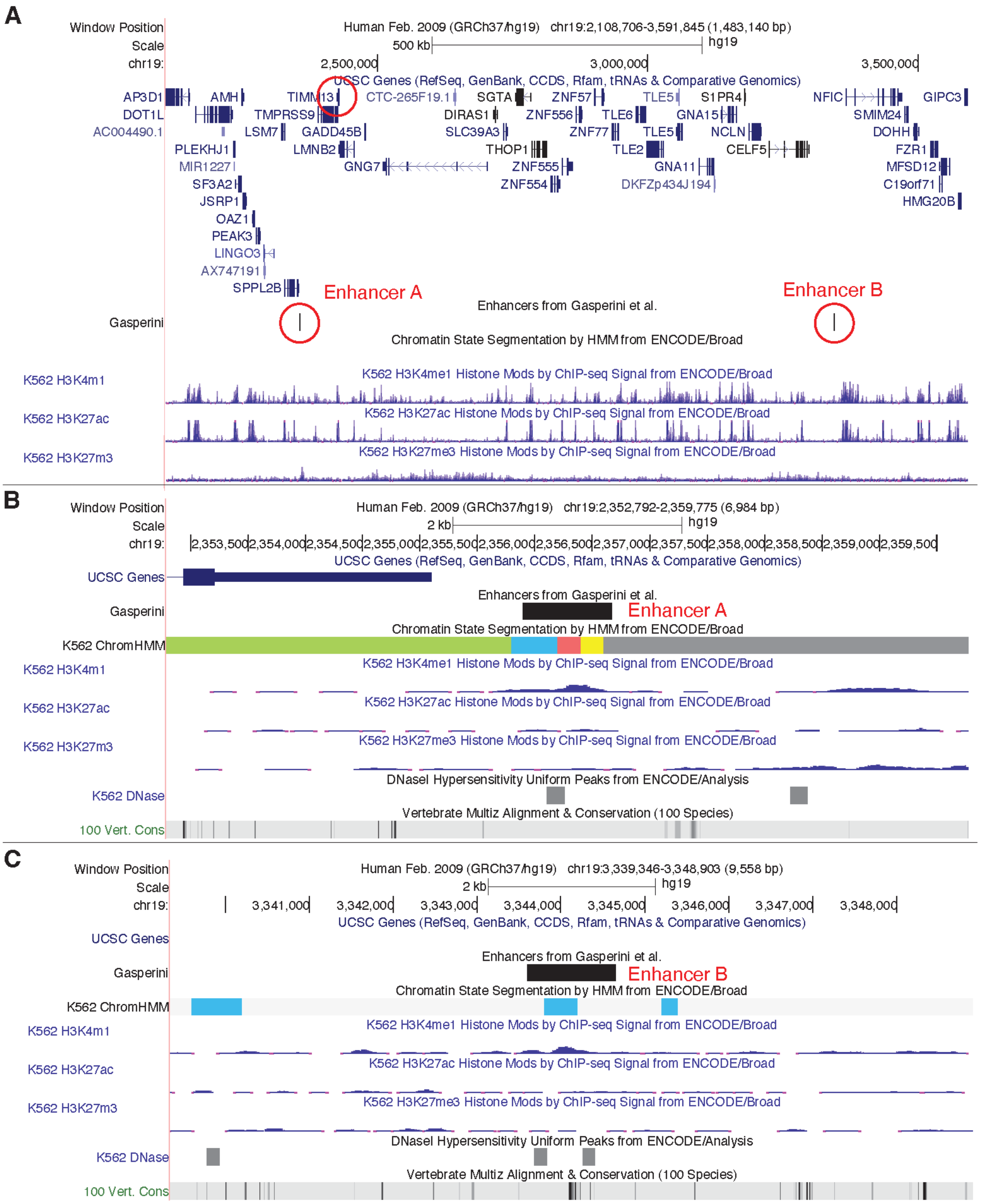


**Figure S5. A pair of enhancers with a negative interaction effect on the expression of *TIMM13*. A)** Overview showing entire genome regions and both enhancers. **B)** Zoom-in of enhancer A region. **C)** Zoom-in of enhancer B region.

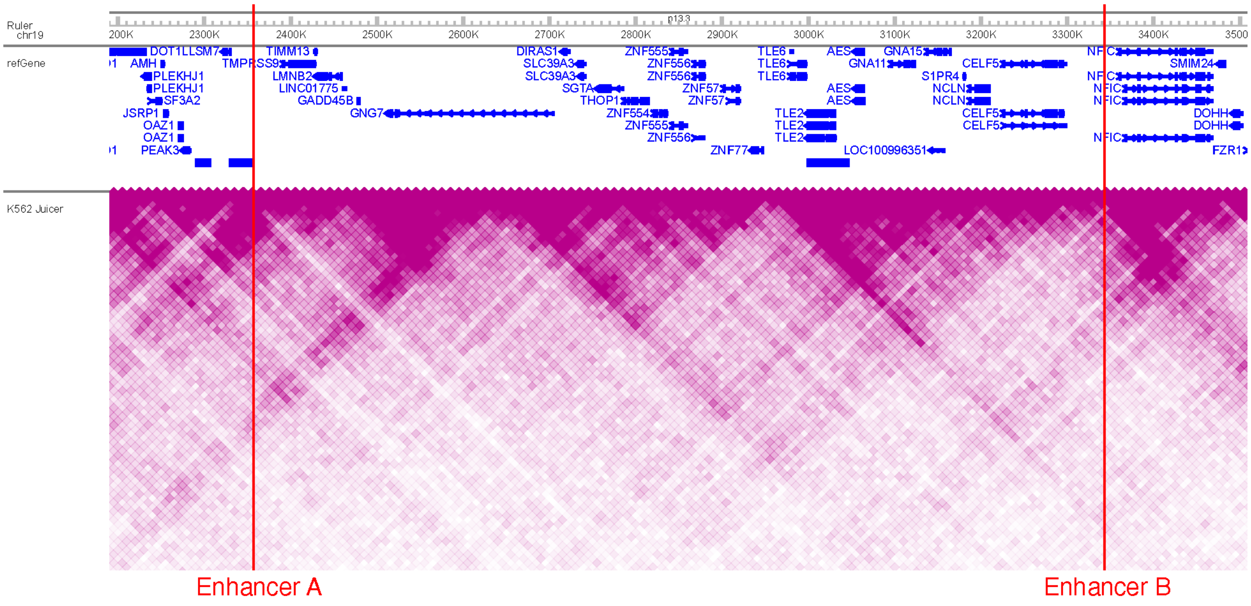


**Figure S6. Enhancers with a negative interaction effect on the expression of TIMM3 are in different topological associating domains (TADs).** The heatmap shows the Hi-C contact frequency in K562 cells[46] from the WashU epigenome browser[47] and the locations of both enhancers.

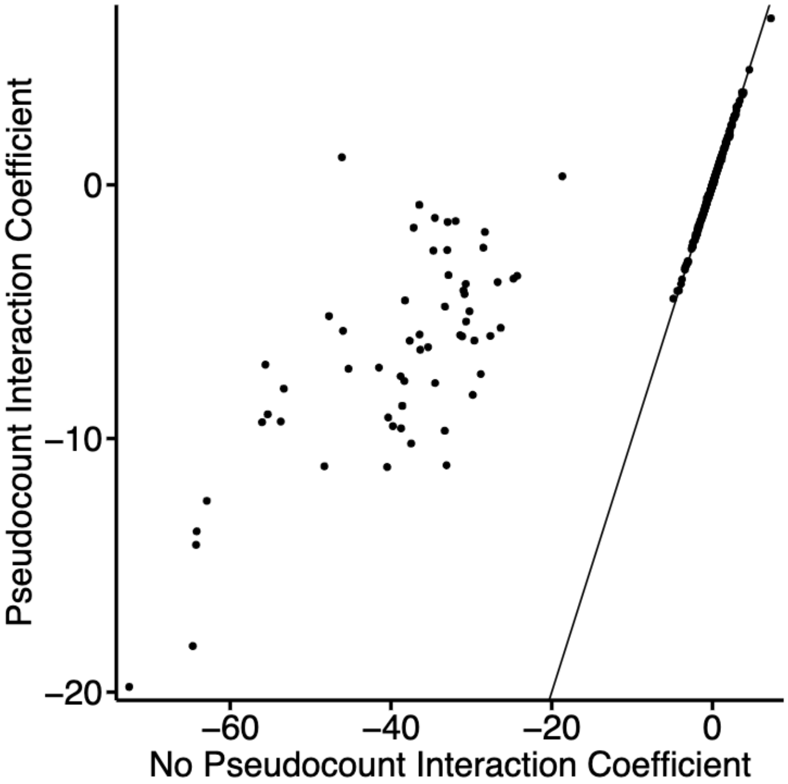


**Figure S7. Outlier interaction coefficient estimates are moderated by introduction of a pseudocount.** Magnitude of interaction term coefficients for 330 enhancer-enhancer pairs when adding vs. not adding a pseudocount of 0.01 to adjust the gene expression. The inclusion of a pseudocount greatly reduces the magnitude of outlier interaction coefficient estimates (note difference in x and y axis scales).

#### Supplementary Tables

**Table S1. Data from *NMU* RT-qPCR experiment.** Provided separately as an Excel file.

**Table S2. Fitting GLiMMIRS-base to simulated data comparing perturbation probability to indicator variable for $X_{perturb}$.** Pearson correlation (Pearson $r$, p_val) between reference and estimated coefficient values for each coefficient in the baseline model when fitting with guide-efficiency derived value of $X_{perturb}$ versus with an indicator (0/1) value for $X_{perturb}$.

| $X_{perturb}$ | **term** | **Pearson $r$** | **p_val** |
|---|---|---|---|
| indicator | (Intercept) | 0.997 | 0.00E+00 |
| indicator | guide.eff | 0.811 | 1.65E-234 |
| indicator | s.score | 0.974 | 0.00E+00 |
| indicator | g2m.score | 0.983 | 0.00E+00 |
| indicator | percent.mito | 0.925 | 0.00E+00 |
| probability | (Intercept) | 0.997 | 0.00E+00 |
| probability | guide.eff | 0.862 | 1.07E-296 |
| probability | s.score | 0.974 | 0.00E+00 |
| probability | g2m.score | 0.983 | 0.00E+00 |
| probability | percent.mito | 0.925 | 0.00E+00 |

**Table S3. Fitting GLiMMIRS-base to simulated data comparing different levels of noise in guide efficiency estimates.** Pearson correlation ($r$), between reference and estimated coefficient values for each coefficient in the baseline model when fitting with perturbation probabilities $X_{perturb}$ calculated from different sets of noisy guide efficiency estimates ($D = 1$ high noise; $D = 10$ moderate noise, $D = 100$ low noise).

| D | term | Pearson $r$ | p_val |
|---|---|---|---|
| 1 | (Intercept) | 0.9970442026 | 0 |
| 1 | guide.eff | 0.3706747091 | 9.85E-34 |
| 1 | s.score | 0.972020145 | 0 |
| 1 | g2m.score | 0.9808390969 | 0 |
| 1 | percent.mito | 0.9031908794 | 0 |
| 10 | (Intercept) | 0.9970706885 | 0 |
| 10 | guide.eff | 0.7807013135 | 5.39E-206 |
| 10 | s.score | 0.9720915922 | 0 |
| 10 | g2m.score | 0.9815630149 | 0 |
| 10 | percent.mito | 0.9034224054 | 0 |
| 100 | (Intercept) | 0.9970696407 | 0 |
| 100 | guide.eff | 0.8451032834 | 1.12E-273 |
| 100 | s.score | 0.9720874449 | 0 |
| 100 | g2m.score | 0.9815627591 | 0 |
| 100 | percent.mito | 0.903438156 | 0 |

**Table S4. Results from applying GLiMMIRS-base to high-confidence enhancers from Gasperini et al.** (Provided separately as .csv file).

**Table S5. Power analysis for GLiMMIRS-int.** Power analysis results for different values of $\lambda$ (lambda) and interaction effect sizes (effect.size). True positive rate (TPR) is calculated from simulated data for interacting enhancer pairs and false positive rate (FPR) is calculated from simulated data for non-interacting enhancer pairs. (Provided separately as .csv file).

**Table S6. Results from applying GLiMMIRS-int to high-confidence enhancers from Gasperini et al.** (Provided separately as .csv file).

**Table S7. Results from applying GLiMMIRS-int to complete set of testable enhancers from Gasperini et al.** (Provided separately as .csv file).

**Table S8. Summary of Cook's distance estimates.** Cook's distance for the interaction coefficient was computed for each cell with joint perturbations for the 31 enhancer-pairs with significant interaction effects. (Provided separately as .csv file).

**Table S9. Results from applying GLiMMIRS-base to candidate enhancers for *PTPRC* using data from Morris et al. 2023** (Provided separately as .csv file).

**Table S10. Results from applying GLiMMIRS-int to candidate enhancer pairs and *PTPRC* using data from Morris et al. 2023.** (Provided separately as .csv file).

**Table S11. Enformer gene expression predictions in the presence of perturbations to individual enhancers and joint perturbations for 2136 enhancer pairs and corresponding target genes tested.** (Provided separately as .csv file).

**Table S12. Results from applying GuideScan 2.0 to Gasperini et al. guide sequences.** (Provided separately as .csv file).

**Table S13. Ensembl to HGNC gene mapping used for cell cycle analysis.** (Provided separately as .csv file).

**Table S14. Cell cycle S scores computed with Seurat.** (Provided separately as .csv file).

**Table S15. Cell cycle G2M scores computed with Seurat.** (Provided separately as .csv file).
